# Single-virus fusion measurements reveal multiple mechanistically equivalent pathways for SARS-CoV-2 entry

**DOI:** 10.1101/2021.05.04.442634

**Authors:** Anjali Sengar, Marcos Cervantes, Sai T. Bondalapati, Tobin Hess, Peter M. Kasson

## Abstract

SARS-CoV-2 binds to cell-surface receptors and is activated for membrane fusion and cell entry via proteolytic cleavage. Phenomenological data have shown that SARS-CoV-2 can be activated for entry at either the cell surface or in endosomes, but the relative roles in different cell types and mechanisms of entry have been debated. Here we use single-virus fusion experiments and exogenously controlled proteases to probe activation directly. We find that plasma membrane and an appropriate protease are sufficient to support SARS-CoV-2 pseudovirus fusion. Furthermore, fusion kinetics of SARS-CoV-2 pseudoviruses are indistinguishable no matter which of a broad range of proteases was used to activate the virus. This suggests that fusion mechanism is insensitive to protease identity or even whether activation occurs before or after receptor binding. These data support a model for opportunistic fusion by SARS-CoV-2, where subcellular location of entry likely depends on the differential activity of airway, cell-surface, and endosomal proteases, but all support infection. Inhibiting any single host protease may thus reduce infection in some cells but may be less clinically robust.

**Importance:** SARS-CoV-2 can use multiple pathways to infect cells, as demonstrated recently when new viral variants switched dominant infection pathways. Here, we use single-virus fusion experiments together with biochemical reconstitution to show that these multiple pathways coexist simultaneously and specifically that the virus can be activated by different proteases in different cellular compartments with mechanistically identical effect. The consequences of this are that the virus is evolutionarily plastic and that therapies targeting viral entry should address multiple pathways at once to achieve optimal clinical effects.

## Introduction

The SARS-CoV-2 betacoronavirus has spread globally since late 2019, causing over 600 million confirmed infections and over 6.5 million deaths as of November 2022 (1). As with many coronaviruses, it binds to a cell-surface receptor and is activated for membrane fusion and cell entry by proteolytic cleavage of its spike protein by a host protease. The primary receptor for SARS-CoV-2 binding, as with SARS-CoV binding, is ACE2 (2-6). This is a common, but not universal, binding receptor among betacoronaviruses infecting humans (7); MERS-CoV utilizes DPP4 as a means of host-cell attachment (8). However, the biochemical steps following viral attachment and their roles in cell entry and physiological infection vary more substantially. Canonical descriptions of viral glycoprotein-mediated fusion typically use the terms “priming” for proteolysis that leaves the glycoprotein in a fusion-competent form and “triggering” for receptor-binding or protonation events that trigger a fusogenic conformational change. As we will discuss, SARS-CoV-2 has at least two proteolysis events, one cleaving spike (S) to form S1 and S2, and one that cleaves S2 to form a fusion-active S2’ fragment. The sequencing of the final proteolysis and receptor binding may be flexible, as our data suggest. We therefore will follow an alternate convention in the literature to use “activation” for the second proteolysis step (9).

The host protease responsible for activation and the subsequent subcellular location of viral membrane fusion vary substantially among betacoronaviruses (10). SARS-CoV was canonically thought to utilize cathepsins present in late endosomes for activation, entering via the endocytic pathway (11-13), although it can also enter at the cell surface (14, 15) in a TMPRSS2-dependent fashion, likely including in respiratory epithelial tissues (16). MERS-CoV can be activated by TMPRSS2 (17, 18), permitting cell-surface entry, although infectivity requires two proteolytic steps, the first of which may be furin-mediated (19-21). Initial data on SARS-CoV-2 have indicated the potential for each of these activation and entry mechanisms (6, 22-27), with some ensuing debate about the biochemical and physiological relevance of cell-surface versus endosomal entry and whether this depends on cell type. In particular, variations in the efficiency by which SARS-CoV-2 variants of concern infect lung parenchyma, bronchi, and nasal epithelial tissues have been correlated with differential protease expression, subsequent transmissibility, and severity of disease (28-30).

The site of host proteolytic cleavage limits how early a virus can undergo fusion and entry, but it does not by itself establish the site of entry. A second trigger may be required, as in the case of Ebola (11, 31, 32), the subcellular compartment where proteolysis occurs may not be permissive for entry. Targeted proteolytic inhibitors and inhibitors of intracellular trafficking provide information on the sites of entry (6, 33), but they also potentially perturb membrane composition. An alternative is to biochemically isolate subcellular compartments and exogenously trigger fusion (34). This permits more biochemically precise and controlled testing of cellular requirements for viral entry.

Studying viral entry in reconstituted systems has yielded substantial insight into the biochemical requirements for entry, including the effects of membrane composition and receptor chemical structure (34-42). The viral membrane itself does appear important to mechanism, whether via distinct composition or distinct spike-protein organization, as both coronavirus and influenza virus cell-cell fusion have well-described differences from virus-cell fusion (43-45). Single-virus assays have enabled an added level of mechanistic sophistication, as they permit analysis of heterogeneity among viral particles and straightforward estimates of stoichiometry from single-event waiting times (36, 38, 40, 46-48). As a complement to biochemical reconstitution, single-virus fusion studies on isolated cellular membranes held under exogenous control (34, 36, 49) permit both the control of reconstituted systems and the physiological membrane environment of cells. This is the approach reported here.

Viral membrane fusion is a mechanistically complex process, involving multiple steps of activation and rearrangement of both the viral fusion proteins and the interacting membranes (50-52). Under normal conditions, the activation and rearrangement of the fusion proteins is believed to be rate-limiting for viruses where this has been studied in detail (35, 40, 53, 54). One key mechanistic parameter is the apparent stoichiometry, or the number of viral fusion proteins required to achieve fusion. This is important because if we assume that there exists some free-energy barrier to viral membrane fusion and all participating copies of the fusion protein are equivalent, then each protein contributes the same free energy to overcoming the barrier and achieving fusion (55). The required protein stoichiometry thus reports on both the underlying barrier to fusion and the contribution each fusion protein makes—if either of these change, the protein stoichiometry will change accordingly. This is still true if the above assumptions are not strictly correct: should fusion proteins differ in their energetic contribution, the apparent stoichiometry will still reflect any changes to the fusion protein contributions. Stoichiometry can be estimated from single-virus fusion kinetics using a number of techniques (38, 46, 56) and thus provides a key parameter for assessing viral entry mechanisms. Because factors such as virus-to-virus heterogeneity can complicate these estimates (57), a more stringent alternate comparison is to assess the statistical agreement between cumulative distribution functions for fusion, plots where the fraction of particles fused is assessed versus time. Both techniques are used in this study.

Here we report single-virus fusion studies of SARS-CoV-2 pseudovirions with isolated plasma membrane and controlled exogenous protease treatment. We show that SARS-CoV-2 spike protein can be activated for fusion by a diverse range of host proteases present in the extracellular environment, on the cell surface, and within endosomes. We also show that the plasma membrane is permissive for viral entry through the point of membrane fusion and that additional endosomal factors are not required for membrane fusion. This supports an “opportunistic” model of SARS-CoV-2 entry, where protease activation can occur at several different stages of viral transport and can lead to fusion with whichever cellular membrane is present at the time.

## Results

To measure single-virus fusion mediated by SARS-CoV-2 spike proteins, we used an approach previously developed for influenza and Zika virus (48, 58) and highly similar to related assays for HIV and coronavirus fusion (36, 49). Briefly (Fig. 1), target membranes—in this case host cell plasma membrane vesicles—are immobilized in a microfluidic flow cell. Virus is labeled with a lipid-phase dye at self-quenching concentration, protease-activated, added to the flow cell, and allowed to bind and fuse. Fusion is assessed via lipid mixing between the virus and the host-cell membranes, detected as fluorescence dequenching in optical microscopy.

**Figure 1.**
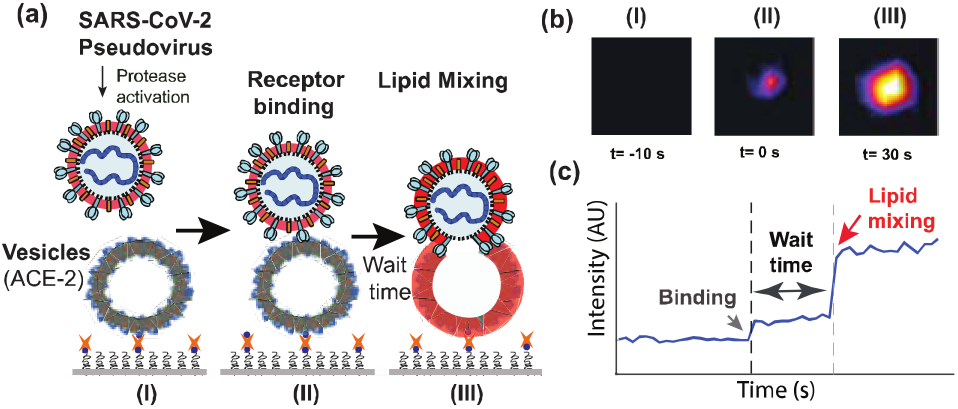
Fusion of SARS-CoV-2 pseudoviruses to plasma membrane vesicles. Panel (A) schematizes the assay design, panel (B) shows fluorescence images corresponding to schematized stages (I-III), and panel (C) shows the corresponding fluorescence intensity trace. Texas Red-DHPE dye is loaded into pseudoviral membranes at a quenching concentration, and plasma membrane vesicles containing ACE2 receptor are immobilized in a microfluidic flow cell (I). Virus-vesicle binding results in a fluorescent spot (II), and lipid mixing between virus and vesicle causes dye dequenching and a further increase in fluorescence (III). This permits measurement of individual pseudovirus fusion events.

In this case, we used three different pseudovirus systems. Pseudoviruses were expressed on HIV, VSV, and MLV backgrounds. All of these pseudoviruses infected Vero cells successfully (Fig. S1) but did not carry SARS-CoV-2 genomes and thus could not replicate. Host-cell plasma membranes were isolated by blebbing giant plasma-membrane vesicles from cultured cells (59), biotinylating these membranes, and then immobilizing them in a flow cell using PLL-PEG-biotin and streptavidin as previously reported (55). Because we observed proteolytic activation of SARS-CoV-2 spike protein to be relatively slow (Fig. S2, Fig. S3), fusion assays were performed by pre-incubating viral particles with the protease of interest and then allowing the activated particles to bind. Cleavage states of the spike protein were assessed via anti-S2 immunoblotting (Fig. S3). In our immunoblots of MLV and HIV as well as previously published immunoblots of the VSV pseudoviruses provided for use in this study (60), a small amount of the spike protein was initially in the S0 form, but the majority was in the cleaved S1/S2 form. Activation of spike protein by further cleavage of S2 to form a S2’ or similar fragment can be observed in the immunoblots. Additional cleavage products likely result from secondary inactivating cleavage events (3). Secondary cleavage events make it challenging to quantitate fraction of S2’ versus S2, but robust fusion is observed with a substantial fraction of S2 spike remaining, suggesting that a relatively small fraction of S2’ cleaved spikes on a (pseudo)virion may be sufficient to support fusion. We also observed activation and fusion without pre-incubation when membrane-bound proteases are present on the target membranes and proteolysis occurs subsequent to receptor-binding (Fig. 2).

**Figure 2.**
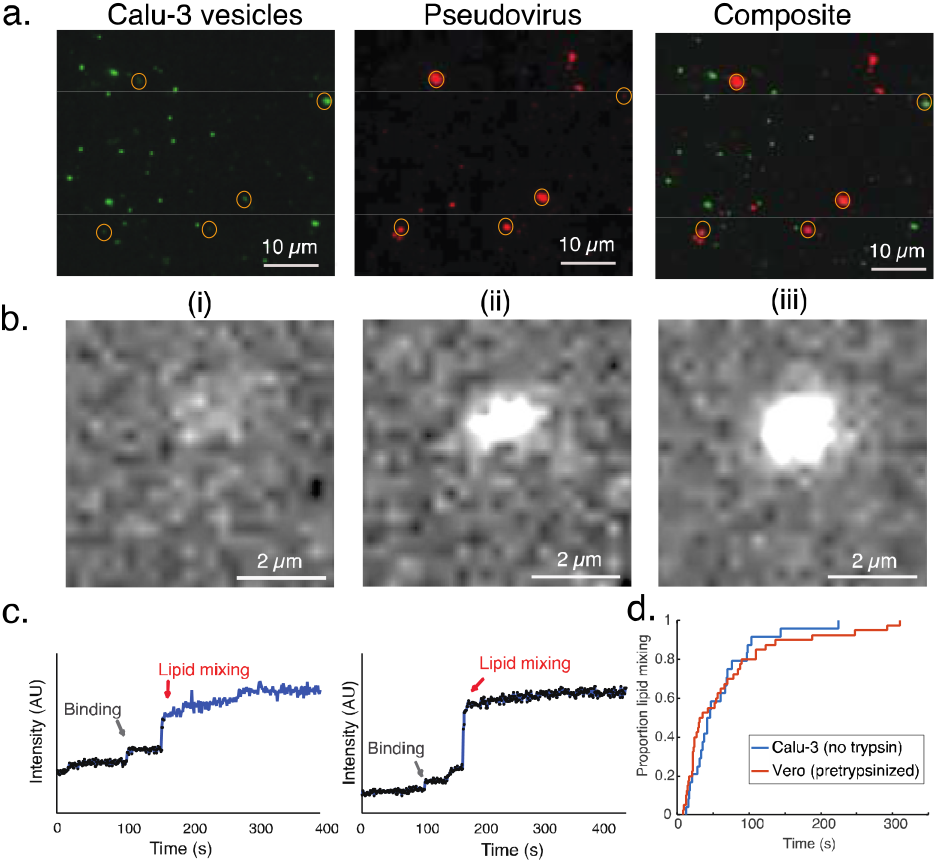
Fusion of SARS-CoV-2 pseudovirus with plasma membrane vesicles in the absence of exogenous protease. Plasma membrane vesicles were obtained from Calu-3 cells and thus contained both ACE2 receptor and TMPRSS2 protease. Cells were labeled with diO membrane dye to facilitate visualization of vesicles, and HIV pseudoviruses were labeled with Texas Red membrane dye. Panel (a) shows fluorescence images of labeled vesicles, pseudovirus, and an overlay showing binding. Multiple co-localized particles are identified via circles in each image. Panel (b) shows a single spot in the Texas Red channel prior to binding (i), at time of binding (ii), and at time of lipid mixing (iii). Panel c) shows two representative fusion traces, plotted as the total intensity of a single-virus spot. Panel (d) shows a cumulative distribution function compiled from the waiting times from binding to lipid mixing measured in 24 lipid-mixing traces similar to (c), compared to a cumulative distribution curve for pre-trypsinized pseudovirus fusing to Vero E6 plasma membrane vesicles. This curve shows the proportion of lipid-mixing events that have occurred by each time on the x-axis.

For testing of exogenous proteases, Vero cell plasma membrane vesicles were used due to their low levels of TMPRSS2 expression yet good ACE2 receptor expression. As a control, plasma membrane vesicles from HEK293T cells that express low levels of ACE2 were immobilized in microfluidic flow cells and HIV pseudovirus allowed to bind in a fashion identical to Vero or Calu-3 plasma membrane vesicles. 21-fold fewer pesudoviruses remained after washing, showing a strong dependence on ACE2 expression (Fig. S4). Furthermore, minimal fusion (< 0.5% of particles dequenching) was observed to Vero cell membranes in the absence of exogenous protease, while more fusion was observed to Calu-3 cell membranes with accompanying higher levels of TMPRSS2 expression (Fig. 2). Prior work has shown that ACE2 mRNA levels are approximately 7-fold higher in Calu-3 cells than in Vero and >10-fold higher than in Vero E6 cells (61).

Analysis of fusion kinetics by proteolytically activated pseudovirions is made challenging by the fact that proteolytic activation and binding are substantially slower than subsequent fusion: for HIV pseudovirions the timescale for activation is 4-5 minutes (Fig. S2) while the timescale for fusion is 71 seconds. We corrected for this by taking virions pre-incubated with protease that then show distinct binding and fusion events on fluorescence microscopy (Fig. 1). These selection criteria may bias the kinetic analysis by not including viruses where binding and fusion occur within the same frame (<1s), but this provides a lower limit for fusion. Additional data on total fluorescence change from lipid mixing integrated across the 133×133 μm microscope field of view are given in the Supplement, and representative fields of view are shown in Fig. S5 and Movies available at doi:10.5281/zenodo.5718787. Cumulative distribution functions were calculated for trypsin-activated fusion by SARS-CoV-2 spike protein on each of the three pseudovirus genetic backgrounds. These results (Fig. 3) show fastest fusion on a VSV background, slowest on an MLV background, with HIV-based pseudovirions intermediate. Single-virus waiting times were statistically different between VSV and either MLV or HIV (p = < 0.001 via 2-tailed Kolmogorov-Smirnov test with Bonferroni correction) but not between HIV and MLV backgrounds (p = 0.69, 2-tailed Kolmogorov-Smirnov test). Furthermore, fusion of HIV pseudovirions to Vero plasma membrane vesicles after pre-incubation with trypsin shows indistinguishable kinetics to fusion of these pseudovirions to Calu-3 plasma membrane vesicles without protease pre-activation (p > 0.5, 2-tailed Kolmogorov-Smirnov test). However, pseudovirions did not fuse to Vero plasma membrane vesicles in the absence of protease treatment. These data suggest that proteolytic activation by TMPRSS2 on Calu-3 cell membranes is not rate-limiting in this case.

**Figure 3.**
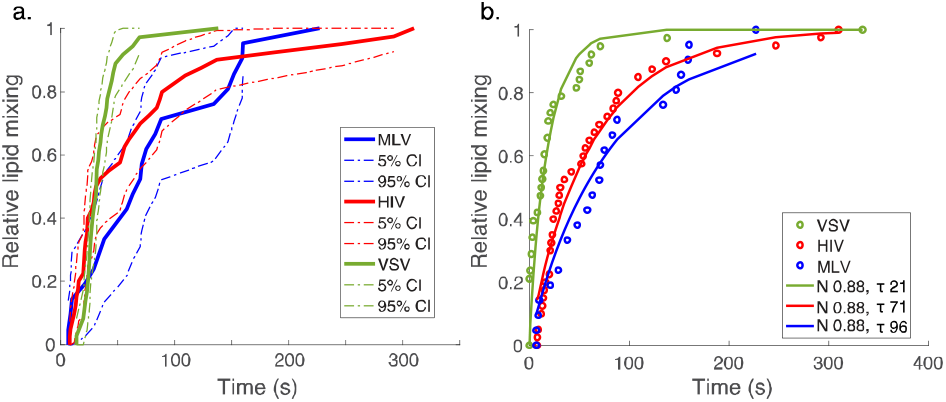
SARS-CoV-2 spike mediates fusion when expressed on different pseudoviral backgrounds. Cumulative distribution functions for single-virus fusion are plotted in panel (A) for SARS-CoV-2 spike protein expressed on VSV (green), HIV (red), and MLV (blue) genetic backgrounds and proteolytically activated using trypsin. All three result in productive fusion (lipid mixing shown here; evidence of downstream fusion shown in Fig. S1). Dashed lines show 90% confidence intervals. Fusion with VSV pseudoviruses is slightly faster than with HIV or MLV pseudoviruses; we hypothesize these differences are due to greater SARS-CoV-2 spike density on the viral surface. Cumulative distribution functions are calculated for 38, 32, and 21 lipid-mixing events for VSV, HIV, and MLV respectively. Fits to these cumulative distribution functions are shown in panel (B) and Fig. S9.

Plasma membrane vesicles are somewhat heterogeneous in size. We took advantage of this to determine the relationship between target membrane curvature and fusion kinetics. Prior work (62) has established that the total fluorescence intensity of a membrane-incorporated dye is proportional to vesicle surface area, so the square root of this intensity is proportional to vesicle radius. Labeling of Vero cell plasma membranes with DiO prior to vesicle production and measurement of lipid-mixing kinetics yielded an assessment of this relationship on a single-vesicle, single-virus level as we have done previously for influenza fusion (63). These measurements showed no correlation (Spearman rank correlation 0.27) between plasma membrane vesicle curvature and time to lipid mixing (Fig. 4).

**Figure 4.**
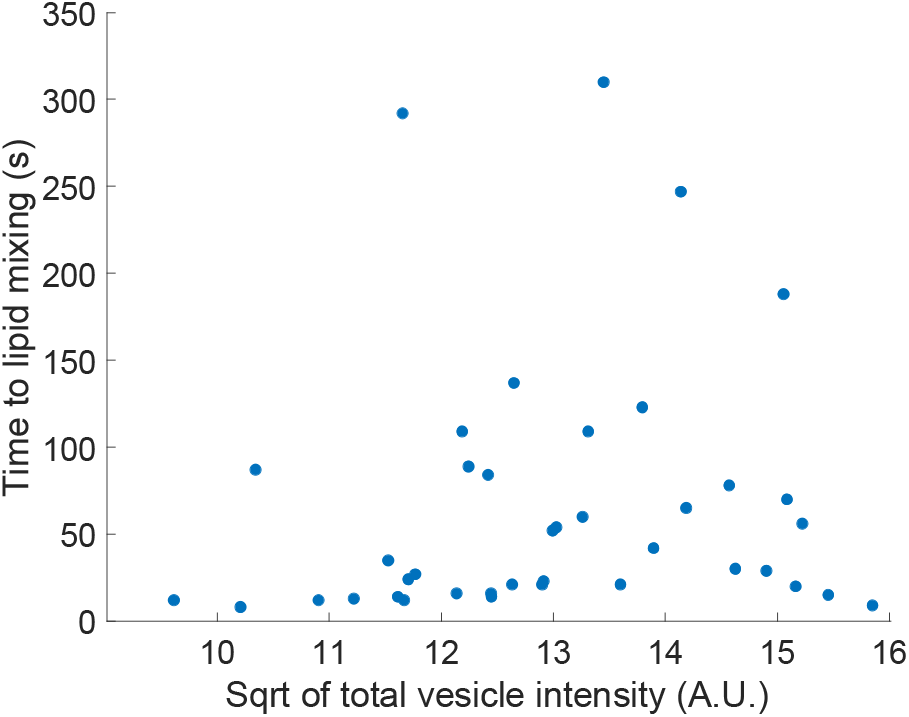
Lipid mixing times are not correlated with plasma membrane vesicle curvature. Plasma membrane vesicle size was assessed by labeling Vero cell plasma membranes with DiO prior to vesicle formation. The total fluorescence intensity of the vesicle in an epifluorescence image is proportional to vesicle surface area, so the square root of total intensity is proportional to vesicle radius. Therefore square-root intensity values are plotted against the time to fusion for individual HIV pseudoviruses activated with 50 μg/mL trypsin and then incubated with the labeled plasma membrane vesicles. No significant correlation was detected (Spearman rank correlation 0.27).

Additionally, to verify that endosomal maturation is not required for productive infection, we incubated Vero cells with 1) SARS-CoV-2 virus-like particles or 2) trypsin-activated SARS-CoV-2 virus-like particles in the presence of 5 nM bafilomycin A, a concentration previously shown to inhibit endosomal entry of SARS-CoV-2 pseudovirus into Vero cells with minimal cytotoxicity (64). As assessed by bulk luciferase assays (Fig. 5, Fig. S6), trypsin-treated particles underwent some inactivation, but the activated ones became independent of bafilomycin A inhibition in TMPRSS2^low^ Vero E6 cells. The majority of trypsin activation occurred prior to ACE2 binding, as demonstrated when the serine protease inhibitor aprotinin was added to trypsinized pseudovirus prior to incubation with cells. This demonstrates that productive virus-like-particle infection does not necessarily require endosomal acidification or the activity of late-endosomal proteases. Our data are highly concordant with the results previously reported using infectious SARS-CoV-2, where pre-treatment of virus with trypsin overcame the inhibitory effects of blocking endosomal acidification (65). Differing results have been reported on this in the literature, including a demonstration using single-virus tracking that inhibition of endocytosis could arrest infection by VSV pseudoviruses (66). The subtle differences between experimental conditions used in these and the single-virus tracking experiments may yield future insight into SARS-CoV-2 biology: some of these may include exposure to protease prior to ACE2 receptor binding, potential alterations in plasma membrane composition and organization induced by dynasore treatment (67) to inhibit endocytosis in the single-virus tracking study, or differences among the viral and pseudoviral constructs used.

**Figure 5.**
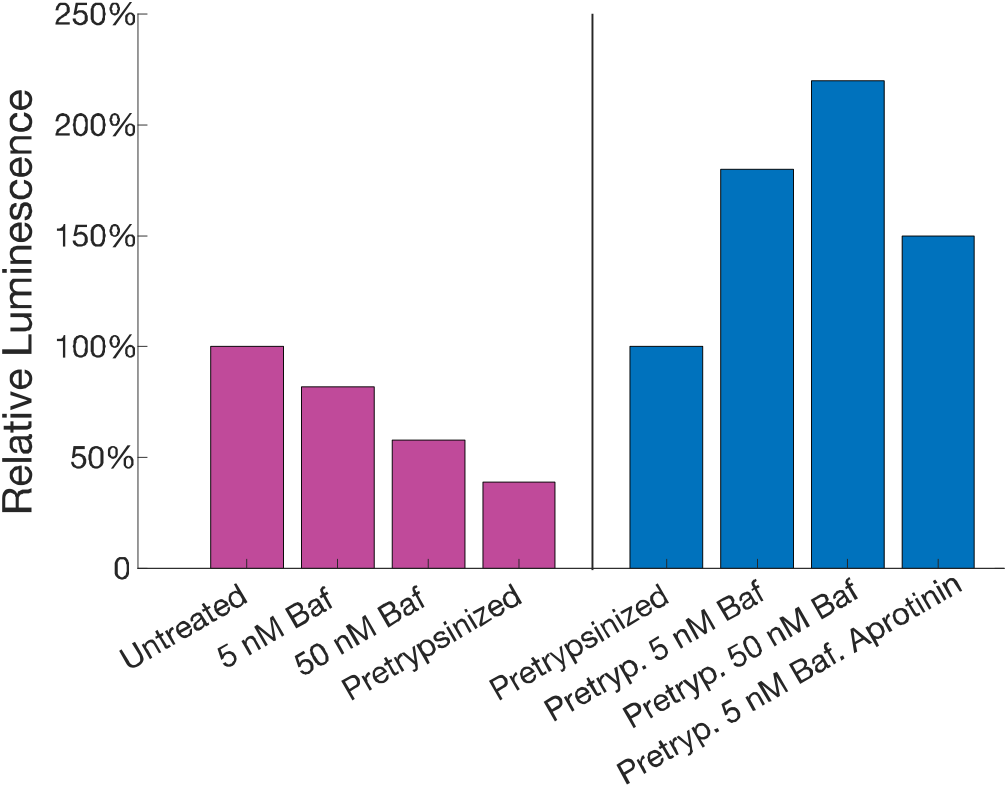
Bafilomycin decreases entry of untreated but not trypsin-activated virus in Vero cells. Pseudoviral entry into Vero E6 cells, which express low levels of TMPRSS2, was assessed in the presence of varying levels of the V1 ATPase inhibitor bafilomycin, which prevents endosomal acidification. Untreated viral entry was inhibited by bafilomycin but not trypsin-treated virus, suggesting that proteolyzed spike proteins permit entry independent of endosomal acidification. Trypsin treatment itself causes a modest inactivation performed long enough but also activates spike proteins. In the final condition, the serine protease inhibitor aprotinin was added after trypsin treatment but prior to incubation with cells in the presence of 5 nM bafilomycin. The resulting luciferase signal was decreased only 17% relative to pretrypsinized virus entering cells in the presence of 5 nM bafilomycin, suggesting that most trypsin activation occurs prior to ACE2 receptor binding.

Because pseudoviruses have a different budding and assembly process from infectious SARS-CoV-2, we also compared single-virus lipid mixing kinetics of virus-like particles (VLPs) formed using S, M, N, and E proteins from SARS-CoV-2 (68). Such particles undergo an assembly process similar to infectious virus and bud from the endoplasmic reticulum (69) Single-virus lipid-mixing kinetics by SARS-CoV-2 VLPs (Fig. S7) was slightly slower but not statistically different from MLV and HIV pseudovirus fusion; VSV pseudoviruses underwent lipid mixing significantly faster than all three other types of particles (p < 0.001, Kolmogorov-Smirnov test with Bonferroni correction).

In most viral fusion systems studied thus far, lipid mixing has been best described as a stochastic process requiring N independent kinetic steps, where each independent step represents the activation of one fusion protein (38, 40, 55, 70, 71). The minimum number of fusion proteins in the most-likely pathway for fusion can then be calculated using the inverse of the normalized variance for the single-event fusion time: N_min_ = <T>^2^ / var(T) for lipid-mixing times T (57, 72). The apparent fusion protein stoichiometry for SARS-CoV-2 spike fusion is between 1 and 3: N_min_ values were calculated at 0.89 (95% CI 0.61-1.6) for HIV pseudoviruses, 1.8 (95% CI 1.2-3.4) for MLV pseudoviruses, and 0.29 (95% CI 0.12-0.84) for VSV pseudoviruses. Interestingly, both HIV and VSV pseudoviruses show N_min_ values of less than 1. This has previously been discussed in the context of *dynamic disorder* or fluctuations in single-molecule reaction rates (46, 57, 73) or as a consequence of non-linear reaction pathways (72, 74). In this case, we hypothesize that the observations are due to *static disorder* or heterogeneity: multiple populations of pseudovirus that have different fusion rates due to different spike protein densities or morphologies. Immunostaining of pseudovirions with anti-spike antibodies can explain the difference between MLV and VSV fusion rates (Fig. 6) and qualitatively the difference between MLV and HIV. Figure 6 clearly shows a higher median number of spikes per particle for VSV than MLV; the remaining unexplained factor is what makes HIV fusion kinetics (both N_min_ and kinetic curves in Fig. 3) lie roughly intermediate between VSV and MLV rather than more closely resembling VSV. We hypothesize that the remaining difference is due to differences in pseudovirion morphology and spike distribution on the pseudovirion surface (cryo-EM images in Fig. S8). VSV pseudoviral particles are well characterized as having a more VSV-like morphology than retroviral (HIV or MLV) pseudovirus scaffolds. In addition, the spike densities on VSV particles do appear somewhat higher on micrographs, but we do not regard our cryomicrographs as sufficiently high contrast to be definitive.

**Figure 6.**
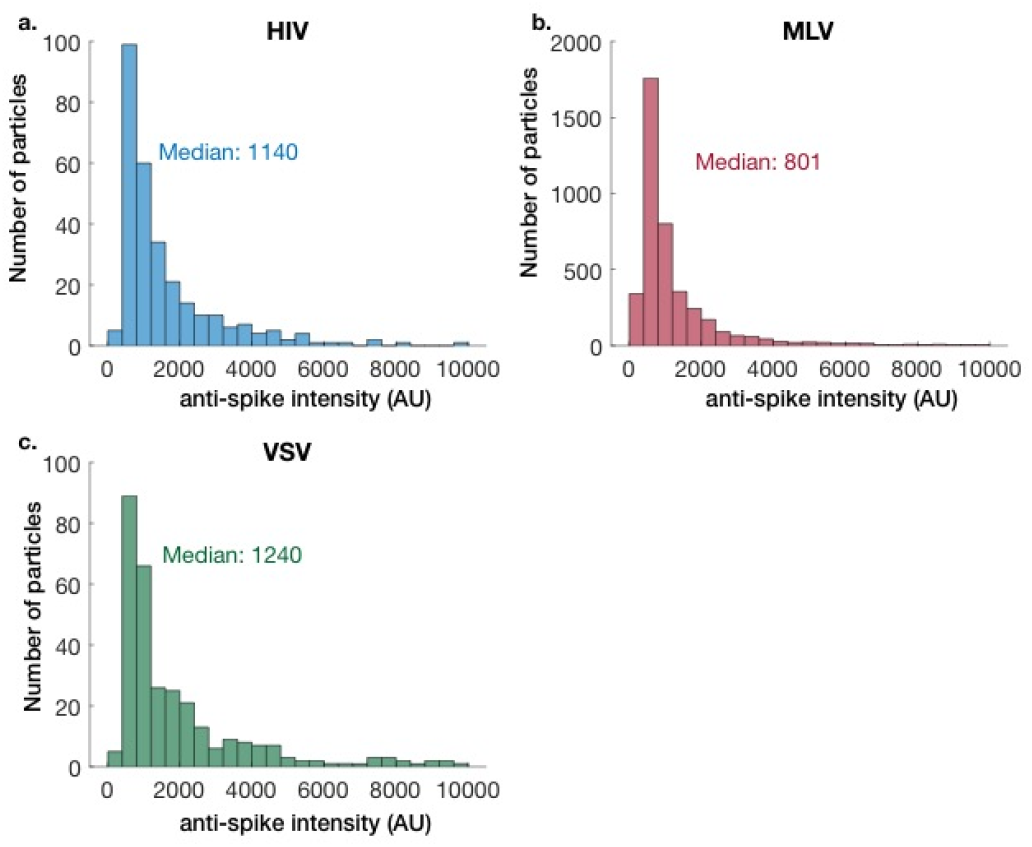
Relative spike protein content on different pseudoviruses. Pseudoviral particles were adhered to a glass surface and probed for SARS-CoV-2 spike protein using indirect immunofluorescence. The fluorescence intensity of each single-virus spot thus is a relative measure of the spike protein content. Histograms compiled from at least three separate experiments per pseudovirus are plotted.

At a coarser level, we hypothesize that the maximum-likelihood stoichiometry is identical between viruses but that the rates of fusion vary across pseudovirus backgrounds, likely due to changes in spike density. We specify maximum-likelihood stoichiometry to allow for stochastic variation in the number of fusion proteins used in individual fusion events but with the hypothesis that the most-utilized stoichiometry is the same across pseudoviruses. We therefore fit cumulative distribution functions to a gamma-distribution model previously used to parameterize fusion waiting times (38), where the fraction of particles undergoing lipid mixing ***f*** is given by 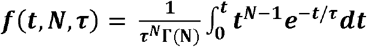, where Γ(N) is a gamma function. We performed two sets of fits, either allowing all parameters to vary independently (Fig. S9) or constraining N to be identical across pseudovirus backgrounds but allowing the parameter ***τ*** to vary (Fig. 3b). The results show high-quality fits with a constraint of common stoichiometry (root-mean-squared error = 0.08, 0.03, 0.06 for VSV, HIV, MLV respectively and Akaike Information Criterion (75) values of 2470 versus of 2530 for unconstrained stoichiometry provide a statistical measure supporting the simpler model of a common stoichiometry), thus suggesting a single most-likely fusion protein stoichiometry. In combination with the single-turnover variance analysis (N_min_) discussed above, we conclude that there is likely static disorder present in the sample i.e. different pseudoviruses have different spike protein arrangements on their surfaces but that the spike protein stoichiometry required for fusion likely does not vary between pseudovirus backgrounds. Instead, the static disorder manifests as different apparent fusion rates. The relative spike protein content of individual virions varies substantially (Fig. 6), supporting this notion.

Fusion mediated by each SARS-CoV-2 spike-bearing pseudovirus was measured at multiple trypsin concentrations to assess differences in fusogenicity of these different constructs. Results show differences in fusion efficiency (Fig. S10) with overall monotonic increases as a function of protease concentration; when efficiency plateaus, we attribute this to saturation. Cumulative distribution functions calculated for HIV pseudovirions treated with 50 μg/mL and 100 μg/mL trypsin show similar fusion kinetics (Fig. 7a): the data are compatible with a model where the stoichiometry of fusion is identical, and the fusion rates are with error of each other. The single-particle waiting time distributions were not significantly different from each other (p > 0.97, 2-tailed Kolmogorov-Smirnov test). These data are again compatible with a similar fusion protein stoichiometry across pseudovirus backgrounds, subject to heterogeneous populations as discussed above; this may however vary between virus-cell fusion and cell-cell fusion as has been suggested for influenza (44). Interestingly, the amount of S2 protein cleaved to form S2’ or a similar activated fragment may differ between proteases (variable band intensity in Fig. S3) but did not appear to alter the kinetics of fusion. Furthermore, the precise molecular weight of these activated fragments varied somewhat between proteases (Fig. S3) but did not alter the measured kinetics.

**Figure 7.**
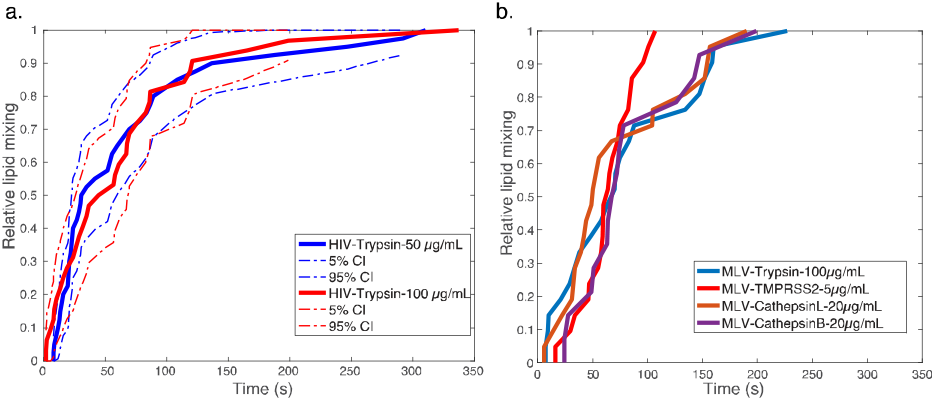
SARS-CoV-2 spike-mediated fusion shows indistinguishable kinetics across protease conditions. Plotted in panel (A) are cumulative distribution functions from single-virus lipid mixing when pseudoviruses were activated with 50 or 100 μg/mL trypsin. Dashed lines are 90% confidence intervals. Plotted in panel (B) are similar cumulative distribution functions with different exogenous proteases (confidence intervals omitted for visual clarity; two-tailed Kolmogorov-Smirnov tests show no significant differences (p > 0.25). Cumulative distribution functions are calculated from 40, 32, 21, 21, 21, and 14 lipid-mixing events for HIV with 50 μg/mL trypsin, HIV with 100 μg/mL trypsin, and MLV with 100 μg/mL trypsin, 5 μg/mL TMPRSS2, 20 μg/mL cathepsin L, and 20 μg/mL cathepsin B respectively.

Finally, we tested the ability of multiple trypsin-like proteases to activate SARS-CoV-2 pseudovirions for fusion. We tested each of the following proteases. Human airway trypsin-like protease (HAT), also known as TMPRSS11D, is a serine protease expressed in both lower and upper airway tissues, found in sputum, and also expressed in other tissues (76-78). HAT has previously been suggested as a candidate for SARS-CoV-2 proteolytic activation (26). Cathepsins B and L are late-endosomal proteases implicated in activation of other viruses including SARS-CoV (12, 13, 22, 79). No increase in fusion efficiency was observed at pH 5 compared to pH 7.4 despite prior structural evidence suggesting pH-dependent conformational difference in SARS-CoV-2 spike(80); it is possible that these changes may affect binding but not fusion. Finally, TMPRSS2 is a cell-surface protease expressed in type 2 alveolar cells among others that has been implicated in MERS proteolytic activation and is a primary candidate for SARS-CoV-2 proteolytic activation (6, 23). Each of these was indeed capable of cleaving (Fig. S2, Fig. S3) and activating SARS-CoV-2 spike protein for fusion (Fig. 7b; Fig. S10; Fig. S11). None of the single-particle waiting time distributions were significantly different from each other (p > 0.25, 2-tailed Kolmogorov-Smirnov test), although TMPRSS2 shows slightly but not significantly faster fusion.

## Discussion

These data support an opportunistic model of SARS-CoV-2 activation and entry, where the virus can robustly utilize multiple parallel pathways for infection with mechanistic indifference. Proteolytic activation by cleavage at the S2’ site can occur in the airway extracellular milieu, at the cell surface, or within late endosomes. This is consistent with prior reports of multiple proteases being capable of activating SARS-CoV-2 (65, 81-84). Furthermore, proteolytic activation can either precede or follow receptor binding; these events need not occur in sequence, and the resulting fusion is mechanistically indistinguishable. We also demonstrate that the plasma membrane is capable of supporting SARS-CoV-2 fusion in addition to the previously reported endosomal fusion site (84, 85). Strikingly, these parallel pathways not only coexist but involve the same number of kinetic steps and thus likely the same mechanism, as probed by single-virus kinetics. We therefore propose a model (Fig. 8) where the site of SARS-CoV-2 entry is determined based on the proteases present in a given tissue and the rates of spike protein cleavage relative to the rates of viral attachment and endocytosis. Where extracellular or cell-surface proteolysis is rapid, entry will tend to occur at the cell surface, whereas if extracellular and cell-surface proteolysis are slow, entry will tend to occur within endosomes. This model is also supported by other cell-biological results (65, 81, 86) that have been published recently (many since initial posting of this manuscript), which we here demonstrate biochemically.

**Figure 8.**
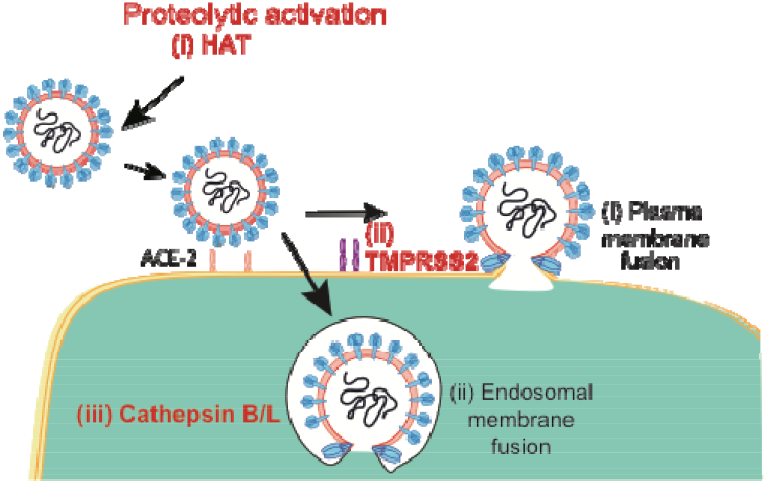
Opportunistic activation and entry of SARS-CoV-2. We propose a model where SARS-CoV-2 undergoes proteolytic activation prior to membrane fusion but where this activation can occur (i) in the extracellular fluid prior to binding (Fig. S11), (ii) at the cell surface after binding (Fig. 2), or (iii) within endosomes (endosomal proteases shown in Fig. 7b). In this model, both (i) plasma membrane and (ii) endosomal membranes are permissive for viral entry. Thus viral entry requires activation and binding to a host membrane, but the requirements for these two are loose rather than stringent.

The key proteolytic triggering event is believed to be formation of the S2’ fragment of the SARS-CoV-2 S protein, which releases the fusion peptide (6, 87) and likely potentiates conformational rearrangement to form the postfusion form of the protein (88). This proteolytic event may be facilitated by conformational changes elsewhere in the spike protein (89, 90). Trypsin, TMPRSS2, and TMPRSS11D are all serine proteases with compatible cleavage sites just upstream of the fusion peptide, forming the canonical S2’ fragment. Cathepsin L, however, has previously been described to cleave SARS-CoV-1 somewhat farther N-terminal to the fusion peptide yet still lead to functional activation (12, 19, 91). Cathepsins have also been described as functionally activating SARS-CoV-2 (33, 82), and our results show that either produce fusion proteins that act with the same kinetics and stoichiometry, suggesting that the additional residues do not impact activation and fusion mechanism. Our results on fusion kinetics thus show that cleavage proximal to the fusion peptide or farther N-terminal are functionally and, we believe, mechanistically equivalent in activating SARS-CoV-2 S for entry.

Our single-virus fusion measurements utilize BSL-2 pseudoviral systems rather than infectious SARS-CoV-2. However, the robustness of the results across multiple pseudovirus backgrounds suggests that the observations are fundamental features of SARS-CoV-2 spike-mediated fusion regardless of the viral core. Furthermore, since observed fusion rates (although not mechanism) do vary with apparent spike protein density, SARS-CoV-2 viral variants that express more functional spike protein are expected to be more infectious. This may be a mechanism for the infectiousness of the D614G mutant (92). We also note that measurements of lipid mixing do not probe viral core exposure and it is unlikely yet possible that differences in protease activation could affect fusion pore opening and viral uncoating. Such a finding would diverge from the body of work on other viral families (40, 93) where activation kinetics primarily affect hemifusion and lipid mixing rather than downstream fusion-pore opening.

The ability of SARS-CoV-2 to utilize parallel entry pathways may contribute to effects where inhibitors targeting a single host protease, such as camostat mesylate, directed at TMPRSS2, are effective in cell and tissue models but show less robust efficacy in clinical trials (6, 94-96). Our data suggest that clinically effective inhibition of proteolytic activation may benefit from combination therapy to target multiple host proteases. Potent monotherapies may sufficiently reduce overall infection to have a clinical effect, but we suggest that multi-targeted therapy may be more efficacious overall. We also show in particular that extracellular proteases such as TMPRSS11D can productively activate SARS-CoV-2 spike for viral membrane fusion, likely in advance of ACE2 receptor binding. Effective protease-targeted therapies thus need to consider the airway extracellular milieu in addition to the cell and tissue types typically used to assess betacoronavirus entry.

The opportunistic activation of SARS-CoV-2 also has important implications for viral evolution and host tissue infection. SARS-CoV-1 was canonically thought of as utilizing the endosomal entry pathway (2, 11-13, 43, 97), although substantial evidence suggests that it, too, can be activated by TMPRSS2 and other proteases and can enter at the cell surface (14, 15, 98, 99). Most recently, while this work was in review, the B.1.1.529 or omicron variant of SARS-CoV-2 was shown to have reduced TMPRSS2 sensitivity and concomitant increased utilization of endosomal entry pathways (29, 85). This change is correlated with differences in relative susceptibility of different airway tissues to infection and changes to resulting pathology. It is thus likely that the ability of SARS-like betacoronaviruses to utilize multiple proteases for entry widens the potential range of tissues that can be infected and facilitates ready adaptation to new host and tissue environments.

## Methods

### Experimental Design

As schematized in Figure 1, plasma membrane vesicles were extracted from either Vero E6 or Calu-3 cells and bound to passivated glass supports in a microfluidic flow cell. Pseudoviral particles expressing SARS-CoV-2 spike protein were fluorescently labeled with Texas Red-DHPE dye at a quenching concentration, either pre-incubated with a specified protease or not, and added to the flow cell. Pseudovirion binding to ACE2 on the plasma membrane vesicle surface was detected as appearance of a dim spot on fluorescence video microscopy, and lipid mixing between the pseudovirion and the plasma membrane vesicle was detected as an abrupt increase in spot intensity. Video micrographs were analyzed to identify bound virions and extract binding-to-fusion waiting times.

The kinetic profiles of these waiting times were then analyzed to compare fusion mechanisms between pesudoviral backgrounds, activation protease, and protease pre-treatment versus activation by membrane-bound TMPRSS2.

### Statistical Analysis

Single-virus waiting time distributions were compared directly between conditions using 2-tailed Kolmogorov-Smirnov tests. Waiting time distributions were also analyzed using a gamma-function model (functional form given in the main text) and via randomness-parameter analysis (46, 57, 74).

### Cell Culture

HEK 293T/17 cells and Vero-E6 cells were cultured at 37 °C and 5% CO_2_ in growth medium containing high glucose Dulbecco’s modified eagle medium (Gibco) supplemented with 10% calf serum (Hyclone), 1% L-glutamine, 1% sodium pyruvate. Calu-3 cells were cultured in minimum essential medium (Gibco) with 12% calf serum (Hyclone), 1% non-essential amino acids, 1% L-glutamine, 1% sodium pyruvate.

### Pseudovirus and virus-like particle preparation

For pseudovirus production, 10 million HEK 293T/17 cells were seeded in a T175 flask 18-24 hours before transfection. Cells at 60% confluence were transfected with 14.7 μg of plasmids with 44.2 μL of Lipofectamine 2000 (Invitrogen) with 14 mL of Opti-MEM. For MLV pseudoviruses, the following plasmids were added at a ratio of 1:1:2: pTG-luc (gift of Judith White), pCMV gag-pol (gift of Judith White), and SARS-CoV-2 spike. Either a full-length SARS-CoV-2 spike construct (BEI NR52310, gift of Florian Krammer) or SARS-CoV2-Δ19 (synthesized *de novo* based on the sequence of Wuhan-Hu-1, codon-optimized spike for mammalian expression, and with the 19 N-terminal amino acids deleted) following previously published protocols (11, 100, 101). For HIV pseudoviruses, the following plasmids, all gifts of Jesse Bloom, were added at a ratio of 1: 0.22: 0.22: 0.22: 0.33: Luciferase-IRES-ZsGreen (BEI NR-52516) : HDM-Hgpm2 (BEI NR-52517) : pRC-CMV-Rev1b (BEI NR-52519) : HDM-tat1b (BEI NR-52518) : Spike-ALAYT (BEI NR-52515) following previously published protocols (102). The same protocol was also scaled down for pseudovirus production in 6-well plates. Transfection media were replaced after 6 hours with 28 mL of growth medium, and supernatant containing pseudovirus particles was collected after 48 hours post transfection, clarified for cell debris at 700 xg for 7 mins at 4 °C and filtered through 0.45μm PES syringe filters. VSV pseudovirus particles expressing full-length SARS-CoV-2 spike were obtained from Behur Lee(103). SARS-CoV-2 virus-like particles were prepared as previously described (68) by co-transfecting plasmids for N (Addgene 177937); M and E (Addgene 177938); and S (D614G N501Y; Addgene 177939) along with a luciferase gene with SARS-CoV-2 packaging sequence PS9 into 293T cells (Addgene 177942). Plasmids were gifts from Jennifer Doudna. For single virus fusion assays, pseudovirus or VLP supernatant was pelleted through 25% sucrose-Hepes-MES (20 mm HEPES, 20 mm MES, 130 mm NaCl (pH 7.4)) cushion at 140,000 x g for 2 h at 4 °C as described previously (100). Pseudovirus pellets were resuspended in Hepes-MES (20 mM Hepes, 20 mM Mes, 130 mM NaCl, pH 7.4) buffer without sucrose to obtain a 100x concentration of the initial volume. Viral aliquots were stored at -80 °C and were thawed no more than once. Pseudoviruses and VLPs were handled under BSL-2 conditions with institutionally approved safety protocols.

### Luciferase Infection assay of pseudovirus

Infection of Vero-E6 cells by MLV and HIV pseudoviruses were performed as described previously (101). Briefly, 125,000 Vero-E6 cells were seeded in a 24-well plate, 18-24 hours before infection. Cell culture media was replaced with OMEM media containing the designated dilution of pseudovirus particles (100x concentrate) adjusted to a total volume of 300 μL per well. Plates were centrifuged for 30 min at 100 xg, 4 °C for pseudovirus binding and then incubated at 37 °C, 5% CO_2_. After 12-18 hours, 200 μL complete media was added to each well. Total pseudovirus protein concentration was determined using a BCA assay (Pierce BCA Protein kit) and used to normalize pseudovirus quantities in luciferase infection assays for comparison of MLV and HIV. Luciferase activity was measured in a plate reader (SpectraMax M5, Molecular Devices) 60-78 hours post infection using the BriteLite reagent (PerkinElmer).

### Plasma Membrane Vesicle Preparation

Vero-E6 and Calu-3 cells natively expressing ACE2 receptors were used for preparation of plasma membrane vesicles following previously published protocols (104). Briefly, 90% confluent cells in a 10-cm dish were labelled with DiO (Invitrogen) for 10 mins. The plate was then washed twice with PBS buffer and twice with a GPMV buffer (10 mm HEPES, 150 mm NaCl, 2 mm CaCl_2_, pH 7.4) to remove the unincorporated label. The cells were then incubated with 5 mL of GPMV buffer containing vesiculating agents (25 mM PFA and 2 mM DTT) for two hours at 37 °C, 5% CO_2_. Plates were then transferred to an orbital shaker for one hour at 37 °C to detach plasma membrane vesicles from the cells. The supernatant was then cleared of cell debris by centrifuging at 100 xg for 10 min and concentrated by centrifuging at 20000 x g for 1 hour at 4 °C. Plasma membrane vesicles were resuspended in a 50 μL GPMV buffer and incubated overnight with 1 μM of biotin-PE solution.

### Fluorescent labelling of pseudovirus particles

For single virus experiments, pseudoviruses were labelled with the fluorescent membrane label Texas Red-DHPE (Invitrogen) following the protocol we have previously used for influenza and Zika viruses (48, 58). Texas Red solution (0.74 mg/mL) in ethanol was mixed with HM buffer (20 mm HEPES, 20 mM MES, 130 mm NaCl (pH 7.4)) at a ratio of 1:40. 50 μL of 100x concentrated purified pseudovirus particles (total viral protein concentration approximately 2-2.5 mg/mL for MLV pseudoviruses as measured via a BCA assay) were mixed with 200 μL of Texas Red-DHPE/HB (HEPES 20 mM, 150 mM NaCl, pH 7.2) buffer suspension and incubated at room temperature in the dark for 2 h on a rocker. After that 2.75 mL of HB buffer was added to the mixture, which was divided into two aliquots. Each aliquot was pelleted at 20,000 x g for 60 mins at 4 °C, resuspended in 25 μL of HM buffer at pH 7.4, stored at 4 °C, and used within one week.

### Protease Treatment

Proteases used were resuspended according to manufacturer protocols: Trypsin-TPCK (Sigma), Cathepsin L (R&D Systems), Cathepsin B (R&D Systems), soluble TMPRSS2 (Abnova), Human Airway Trypsin-like protease (R&D Systems). For each experiment, labeled pseudovirus was diluted 10x in HB with the designated protease concentration and incubated at 37 °C for 1 hour. The mixture was then diluted in 200 μL HB and added to a microfluidic flow cell channel at a rate of 20 μL/min while recording video as described below.

### Microfluidic Flow Cell Preparation

Glass coverslips (24 × 40 mm, No 1.5, VWR International) submerged in a 1:7 solution of 7x detergent (MP Biomedicals) in DI water were heated and stirred for 30 min, until the solution turned clear. Coverslips were then rinsed extensively with DI water, baked in a kiln for 4 hours at 400 °C and left in the furnace overnight for slow cooling. After that, coverslips were rinsed with ultrapure water and sonicated with ethanol in a bath sonicator for 10 min. After a final rinse in ultrapure water, coverslips were dried in the incubator at 100°C, cooled coverslips were then stored in a 50 mL falcon tube and wrapped with parafilm until use.

Microfluidic flow-cell molds were prepared using tape-based soft lithography (105). Briefly, Kapton polyimide tape (Ted Pella) was affixed on a microscopic slide and channels of dimensions (1 mm x 13 mm x 70 μm) were carved using a Cameo cutter-plotter. Excess tape was removed from the channels, and the mold was cleaned using isopropanol before use. Polydimethylsiloxane (PDMS; Sylgard 184) was mixed and poured into mold. After degassing under house vacuum, the PDMS was cured at 60 °C for 4 hours, and flow cells were cut out. Channel inlet and outlet holes were created using a 2 mm biopsy punch.

Glass coverslips were plasma cleaned for 5 mins (Harrick Plasma) before bonding. PDMS flow cells and cleaned glass coverslips were plasma bonded together after plasma activation for 1 min. Immediately, PDMS flow cell channels were coated with 95% PLL-PEG: 5% PLL-PEG-biotin. After 30 min incubation, channels were washed with 1 mL of ultrapure water and 1 mL of Hepes buffer (20 mM Hepes, 150 mM NaCl, pH 7.2). The biotin-PEG layer was functionalized by incubating 15 min with a 0.2 mg/mL solution of neutravidin (Thermo Scientific). Channels were washed with 1mL of HB to remove excess neutravidin. 10 μL of biotin-functionalized plasma membrane vesicles were incorporated into the flow cell channel and incubated overnight at 4 °C for biotin neutravidin binding. Each flow cell channel was washed with 1mL of GPMV buffer to remove unbound vesicles.

### Lipid-Mixing Assay

Each microfluidic flow cell (coated with plasma membrane vesicles) was washed and equilibrated using HB buffer. DiO-labeled vesicles were used for preliminary focus adjustment and flow-cell quality control. To initiate the experiment, 200 μL of protease-treated pseudovirus solution was added to the channel at a flow rate of 20 μl/min, and 1200 video frames were recorded at a rate of 1 frame/s. Single particles were identified using previously developed (106) Matlab filters (see code availability below). For single-particle kinetic analyses, time-trace series data for individual particle ROIs were measured using the time-series analysis plugin in Fiji (107). Single-particle waiting times were calculated as the time between the binding of an individual labeled pseudovirus and fluorescence dequenching.

### Microscopy

Video micrographs were acquired using a Zeiss Axio Observer inverted microscope using a 100X oil immersion objective. A Spectra-X LED Light Engine (Lumencor) was used as an excitation light source with excitation/emission filter sets as follows: Cy5 (Chroma 49009 ET-Cy5), Texas Red (Chroma 49008), DiO/FITC (Chroma 49011). Images were recorded with an Andor Zyla 4.2 sCMOS camera (Andor Technologies) using 16-bit image settings controlled using Micromanager (108) software.

### Cryo EM Imaging

Electron cryo-microscopy imaging and analysis was performed as published previously (34, 109). Briefly, 3 μL of concentrated pseudovirus sample was applied to a carbon-coated grid (2/2-4C C-flats; Electron Microscopy Sciences), blotted with filter paper, and plunge-frozen in liquid ethane. Samples were imaged in a Tecnai F20 Twin transmission electron microscope (FEI, Hillsboro, OR) at -180 °C with a magnification of 29,000x or 50,000x, operating at 120 kV.

### Immunofluorescence

Texas Red labeled pseudovirus particles were bound to DiO labeled target plasma membrane inside a flow cell. After washing excess pseudovirions, the flow cell channel was blocked with 3% BSA in HM buffer for 30 min and then incubated overnight with a 1:100 dilution of anti-Spike primary antibody (BEI NR616, obtained from BEI Resources, NIAID) in 3% BSA at 4 °C. The next day, the flow cell was washed with HM buffer and then infused with a 1:500 dilution of Alexa 647-goat anti-mouse secondary antibody (Invitrogen), incubated for 1h at RT, washed again, and then imaged. DiO, Texas Red and Alexa 647 images were collected sequentially for the same field of view.

### Immunoblots

To assay the activity of different proteases against SARS-CoV-2 S, 8 uL of 100 x concentrated pseudoviral pellets in HEPES-MES buffer were treated with the indicated concentration of protease at 37 °C for 1 h. In addition, time-course experiments were performed for trypsin where HIV pseudovirus particles were incubated with a final concentration of 100 ug/mL Trypsin-TPCK (Sigma-Aldrich) for 1, 10, 20, 60 min at 37 °C. After protease treatment, all samples were then quickly subjected to phenylmethylsulfonyl fluoride (Sigma-Aldrich) treatment at a final concentration of 2 mM for 5 min, 37 °C to stop further proteolysis. Samples were then mixed with 6x Laemmli loading dye with 100 mM final DTT concentration, incubated at 95 °C for 5 min, and loaded on a polyacrylamide gel. Gel electrophoresis was performed using a gradient gel (4-12% Mini Protean TGX gel (Biorad # 4561095)) at 120V for 160 min on ice. Samples were then transferred to PVDF membranes using a wet-electroblotting chamber system (Biorad) in Towbin buffer containing 15% methanol for 90 min at 100V on ice. The membrane was blocked with 5% BSA in TBS containing 0.1% tween-20 (TBS-T) for 1h at 4 °C before incubating with anti-S2 Spike primary antibody (rabbit polyclonal, Sino Biologicals, Cat#: 40590-T62) at 1:1,000 dilution in TBS-T with 5% BSA for 14-16 h at 4°C. After that, the membrane was washed 3 times in TBS-T for 10 min, incubated with labeled secondary antibody (goat anti-rabbit IgG, Abcam ab216777) diluted at 1:8000 in TBS-T with 5% BSA for 2 h at RT, washed again 3 times in TBS-T and imaged using a LI-COR Odyssey gel imager.

## Data availability

Image and time-series analysis code written in Matlab is available on Github at https://github.com/kassonlab/micrograph-spot-analysis as previously reported (106). Data are deposited on Zenodo.

## Author Contributions

Conceptualization: AS, PMK

Methodology: AS, MC, PMK

Investigation: AS, MC, STB, TH

Data Analysis: AS, MC, TH, PMK

Writing—original draft: AS, PMK

Writing—review & editing: AS, MC, STB, TH, PMK

## Acknowledgements

The authors thank Jesse Bloom and Judith White for the gift of reagents and M. Cervantes, G. Morbioli, A. Villamil Giraldo, R. Rawle, and J. White for helpful discussions. Electron cryo-microscopy was performed by K. Dryden at the Molecular Electron Microscopy Core at the University of Virginia. This work was supported by grants from the Commonwealth Health Research Board (207-01-18), UVA Global Infectious Diseases Institute, and Knut and Alice Wallenberg Foundation (KAW2015.0198 and KAW2020.0209) to P.M.K.

