## Supplementary Information for "Single-virus fusion measurements reveal multiple mechanistically equivalent pathways for SARS-CoV-2 entry"

### Figures

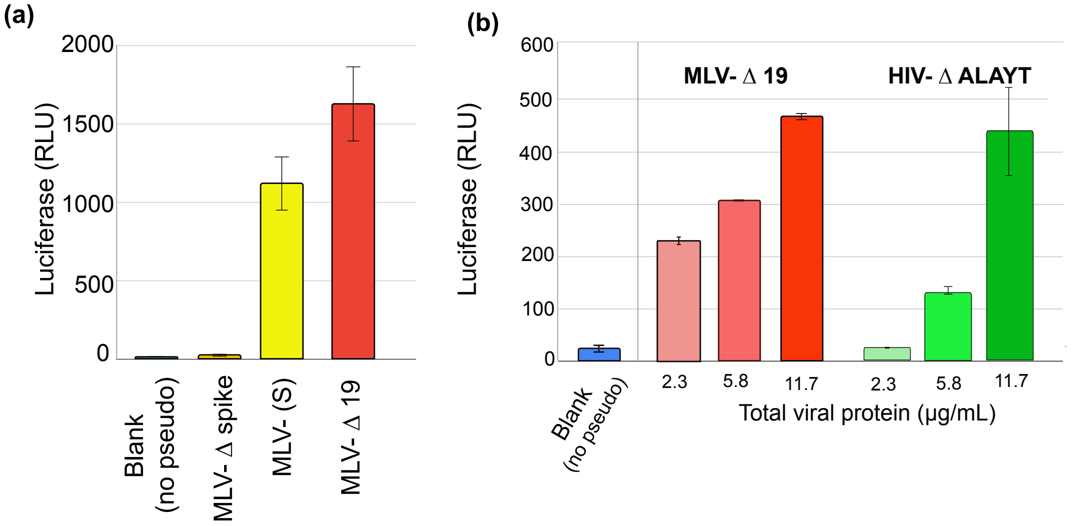


**Figure S1. Productive entry of pseudovirus into Vero E6 cells.** All pseudoviruses contain only a luciferase gene—no other genetic material—so they are not capable of replication. However, an end-to-end measurement of viral entry is luciferase activity in cells, which reflects successful entry and expression of the luciferase gene. As plotted in panel (a), we observe luminescence in Vero E6 cells incubated with SARS-CoV-2 on the MLV pseudovirus background (either full-length or Δ19) or with SARS-CoV-2 on the HIV background (ALAYT cytoplasmic tail mutant (*60*)) but not with MLV pseudoviruses lacking the spike construct or Vero E6 cells alone. Titrations of the MLV Δ19 and HIV ΔALAYT pseudoviruses are plotted in panel (b). Solid bars display mean values of at least three biological replicates, and error bars denote standard deviation. SARS-CoV-2 pseudovirus on the VSV background was a kind gift of Benhur Lee; this reagent was standardized for infectivity, and luciferase assays have been previously published (*61*). Viral quantities are measured in µg/mL total viral protein as estimated by a BCA assay. The ultracentrifugation used to separate viral particles from soluble protein may also capture cellular proteins.

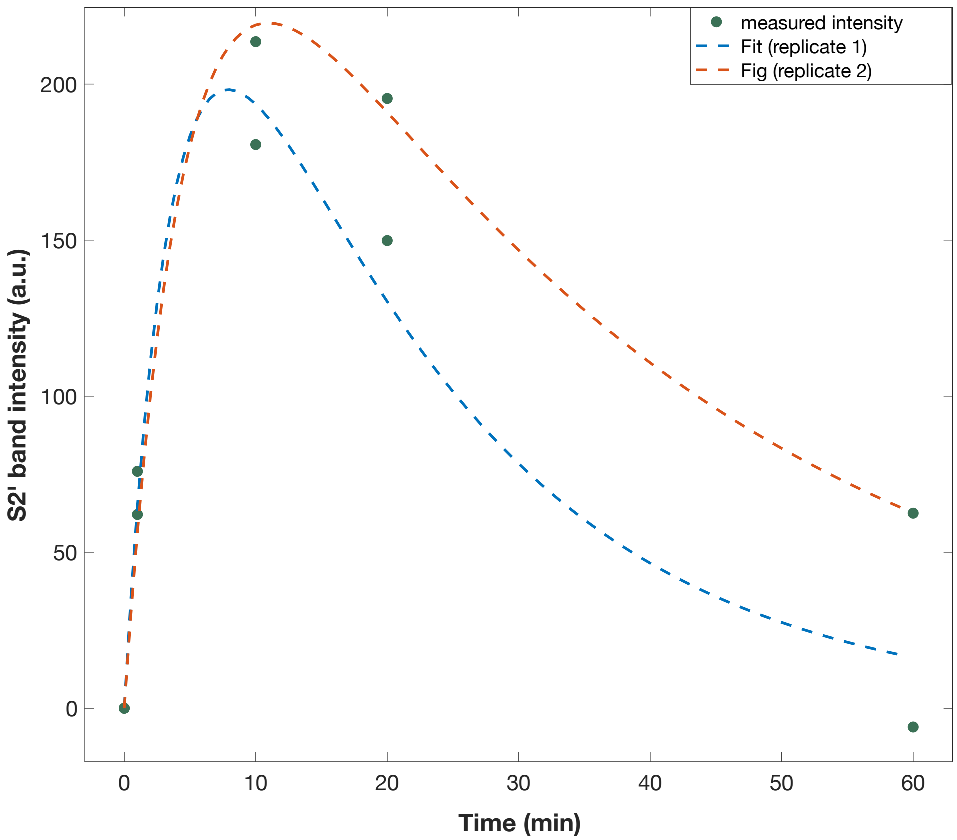


**Figure S2. Cleavage kinetics of SARS-CoV-2 envelope protein by trypsin.** Cleavage was assessed by incubating trypsin with SARS-CoV-2 pseudovirus and then separating cleavage products via SDS-PAGE and immunoblotting with an anti-S2 antibody. The bands corresponding to S2’ and similarly sized cleavage products were measured, and their intensity was plotted versus time and fit using ordinary differential equations: $d[S2']/dt = a * [S2] - b * [S2']$. These fits yielded a timescale (1/a) of 4.0 to 4.8 min for S2 cleavage across two independent biological replicates. Primary immunoblots are shown in Fig. S3. This model assumes that the bands in the 50-64 kDa range include the S2’ cleavage, potentially in addition to other inactivating cleavage events. The fitted kinetics thus represent a lower bound on S2’ kinetics.

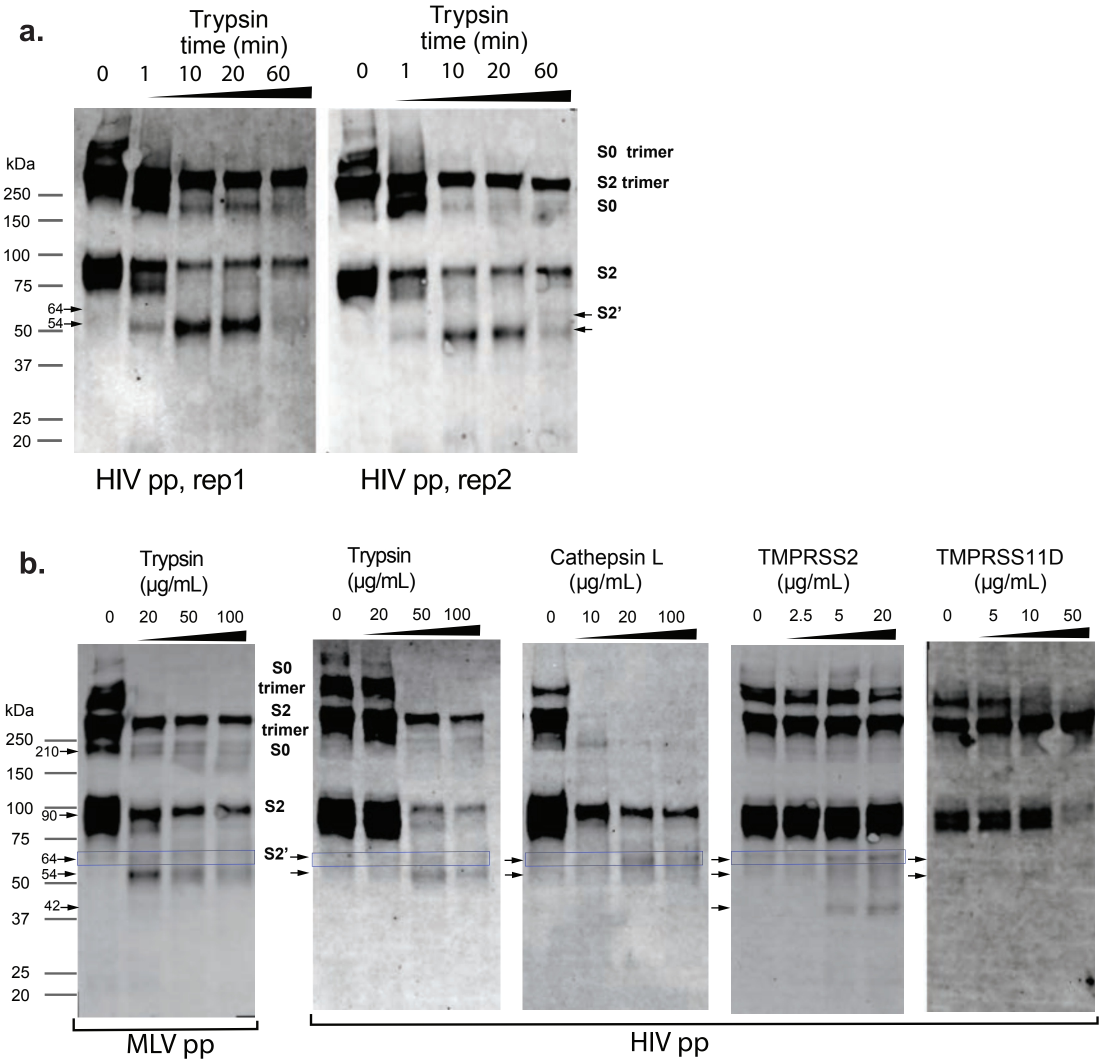


**Figure S3. Cleavage of SARS-CoV-2 envelope protein by different proteases.** Cleavage was assessed by incubating proteases with SARS-CoV-2 pseudovirus and then separating cleavage products via SDS-PAGE and immunoblotting with an anti-S2 antibody. Primary immunoblots are shown above, while results for trypsin cleavage kinetics are quantitated in Fig. S2. Two independent biological replicates (denoted rep1 and rep2) are shown for the trypsin cleavage time-course experiments. Gel band assignment is based on prior reported molecular weights (1, 2) as well as comparison to immunoblots performed using anti-S1 antibodies. The band around 270 kDa likely represents an S2 trimer. Bands are annotated; multiple cleavage products are also observed with many of the proteases, and approximate molecular weights for many of the products are listed as well. The consensus from prior work (3-6) is that the 64 kDa fragment is likely assignable to S2'. The 54-55 kDa fragment could potentially represent either an activating cleavage at a different site (this would be consistent with the band assignment of (7)) or an inactivating cleavage. In the context of a mixed population of spike cleavage forms, as seen in the immunoblots, definitively separating these by fusogenic activity is challenging. But all immunoblots except for TMPRSS11D (HAT) have clearly defined bands at ~64 kDa for the 60-minute time points, which is consistent with the bands assigned to S2' by other prior studies. As noted in the discussion, the majority of the spike protein is in the S1/S2 state without additional protease addition.

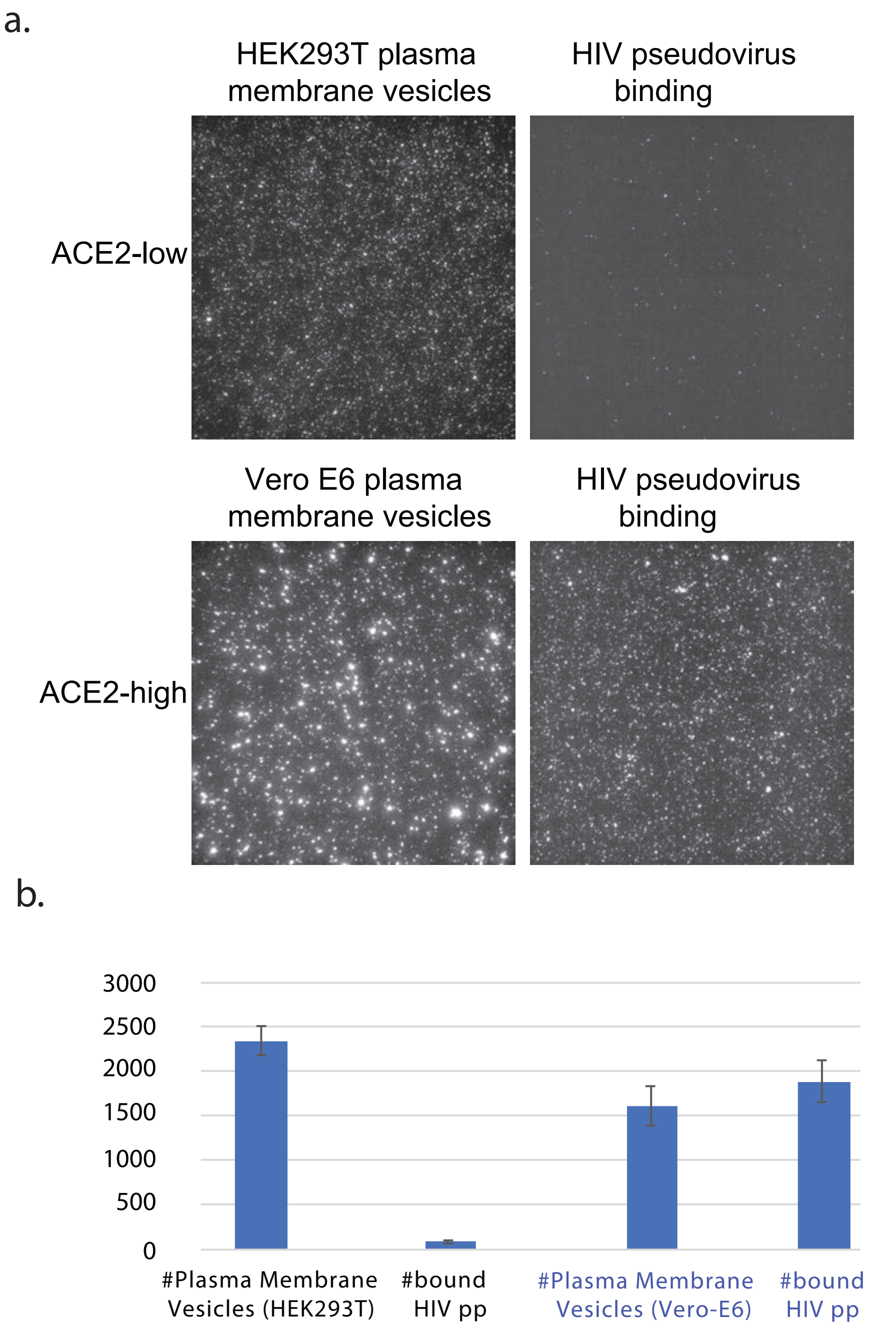


**Figure S4. Reduced pseudovirus binding to HEK293T cells expressing low levels of ACE2.** Plasma membrane vesicles were formed from HEK293T cells and Vero E6 cells and immobilized within microfluidic flow cells. HIV pseudoviruses were allowed to bind, and then unbound virus was washed away and the flow cells imaged both in the DiO channel (left, plasma membrane stain) and the Texas Red channel (right, pseudovirus label). Images in panel (a) are representative fields of view (134 x 134 µm) and are quantified in panel (b) with bootstrapped 95% confidence intervals plotted. Flow cells containing Vero E6 plasma membrane vesicles showed >21-fold higher pseudoviral counts than those containing HEK293T plasma membrane vesicles. Equal amounts of pseudovirus were added to each flow cell.

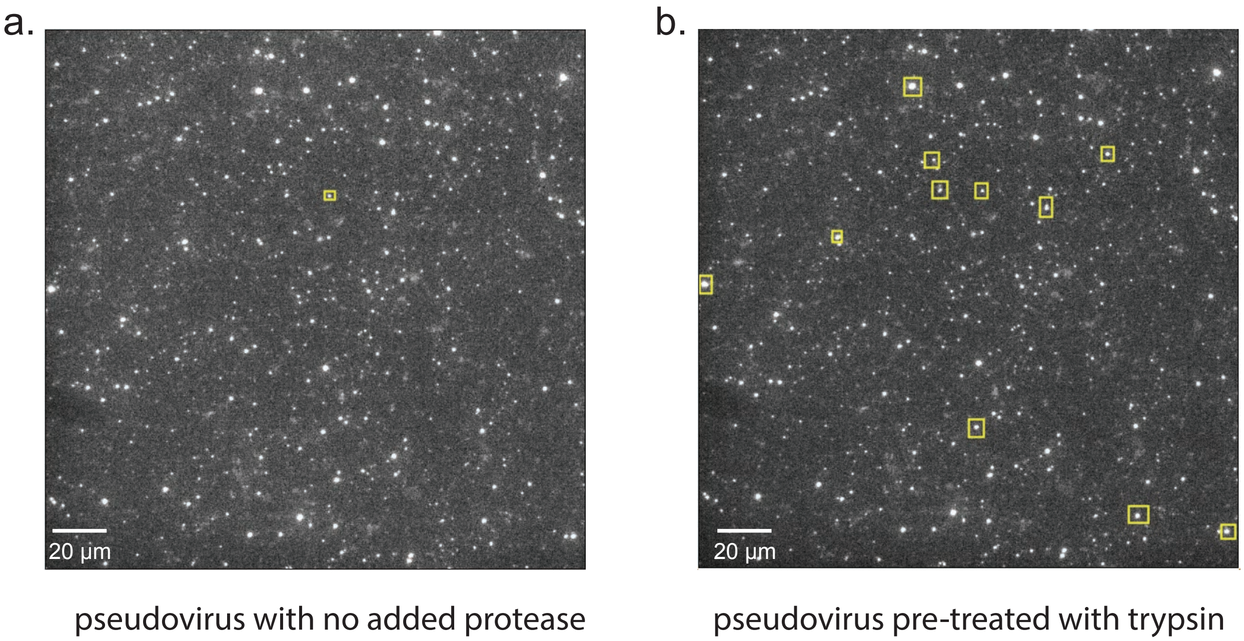


**Figure S5. Minimal fusion in the absence of protease.** MLV pseudoviruses were either pre-treated with trypsin for 60 min or not. These were then added to a flow cell containing Vero-cell-derived plasma membrane vesicles, and video micrographs were recorded to monitor fusion. Representative fields of view are shown for each condition, with spots in the Texas Red channel indicating bound pseudovirus at the end of the experiment. Pseudoviral particles that fused during the experiment are indicated with yellow boxes. Greater than 10-fold more pseudoviruses fused when trypsin was added than when it was not present.

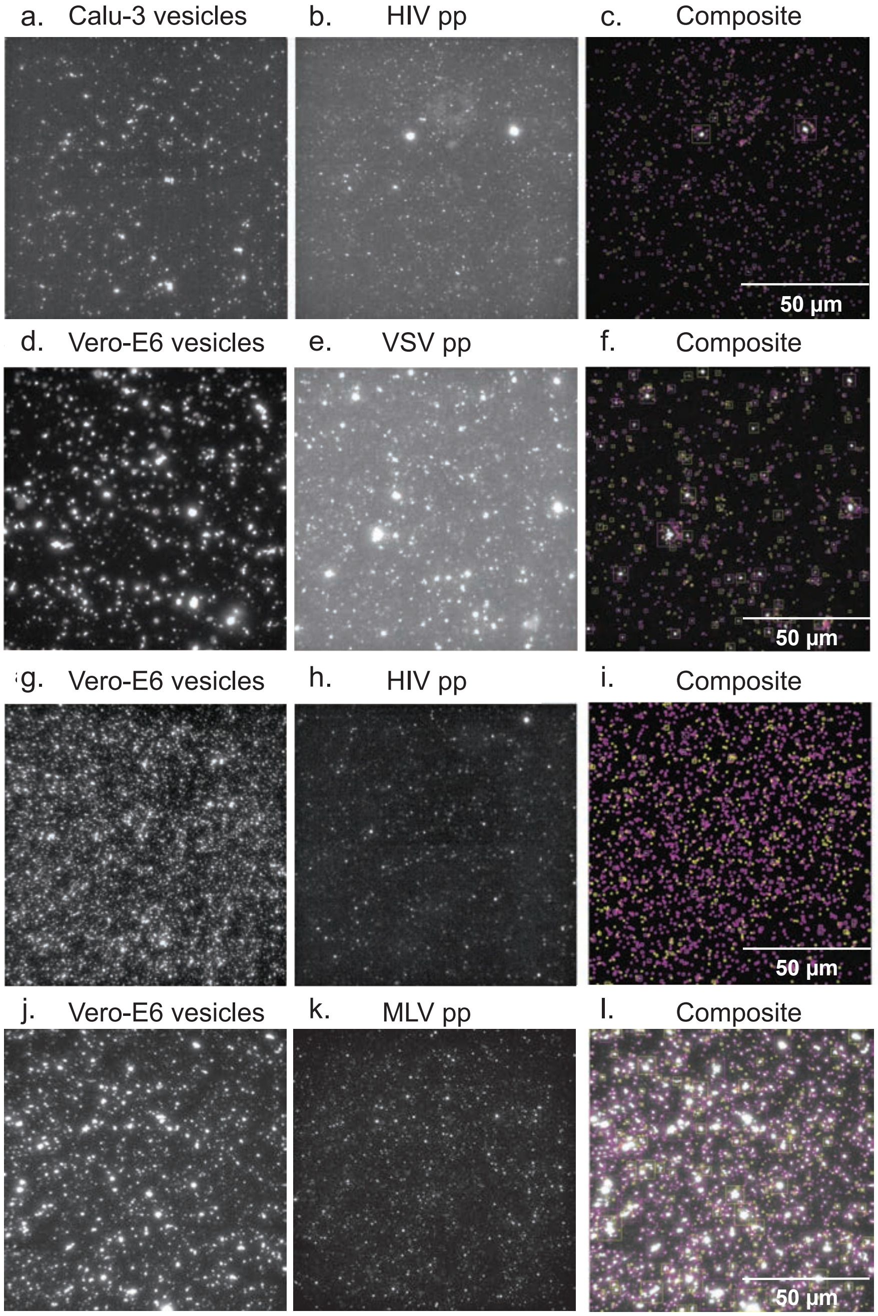


**Figure S6. Representative fields of view for pseudovirus-membrane fusion experiments.** Panels (a)-(l) show representative fields of view for HIV, VSV, and MLV pseudoviruses expressing SARS-CoV-2 S and bound to either Calu-3 or Vero-E6 plasma membrane vesicles. In panel (c), 21% of 699 vesicles had pseudovirus colocalized. In panel (f), 29% of 1969 vesicles had pseudovirus colocalized. In panel (i), 23% of 2288 vesicles had pseudovirus colocalized, and in panel (l), 24% of 1862 vesicles had pseudovirus colocalized. Fusion movies corresponding to each of these experiments are shown in Supplementary Movies 1-4, which are available on Zenodo at https://doi.org/10.5281/zenodo.5718786.

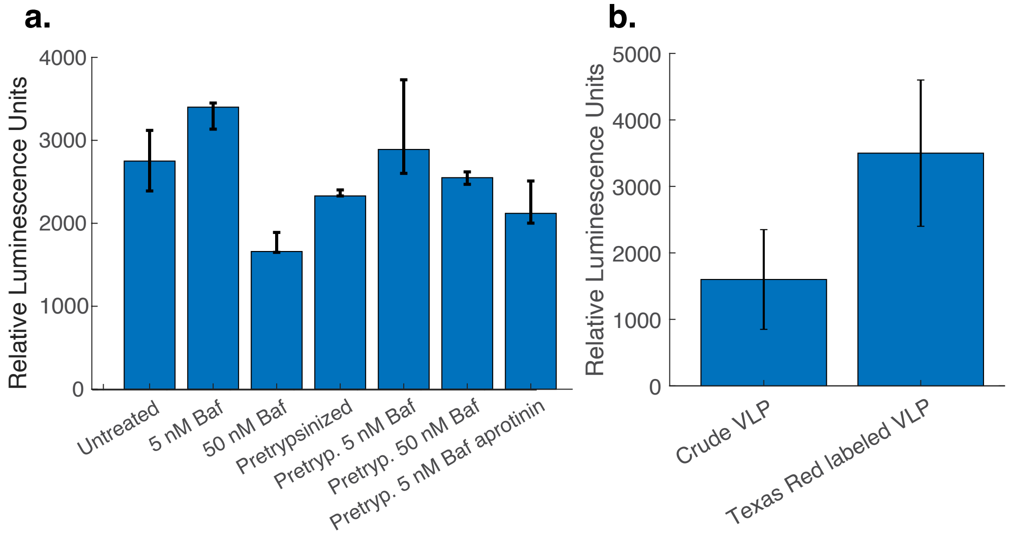


**Figure S7. Effect of fluorescent labeling and endosomal acidification on virus-like particle entry**. Virus-like particles formed by co-expressing S, E, N, and M proteins from SARS-CoV-2 and packaging a luciferase gene were used to infect Vero E6 cells. Panel (a) shows raw luciferase counts corresponding to Figure 5. In addition, pretrypsinized virus in the presence of 30 μM aprotinin and 50 nM bafilomycin yielded 2380 RLU (range 2310-3040 over 3 biological replicates), a decrease of <10% from pretrypsinized virus. Panel (b) shows raw luciferase counts testing the effect of Texas Red labeling on infectivity was tested using this platform. Labeling did not reduce infectivity; viral amount was standardized to total protein concentration, and centrifugation after Texas Red labeling likely removed a noninfectious proteinaceous component, accounting for the increase in luminescence. The experiments in panel (a) did not undergo a centrifugation step. Error bars represent the interquartile range across 5 biological replicas for panel (a) except for the pretrypsinized condition, which had two 4 replicas and the standard deviation across 3 biological replicas for panel (b), and bars denote median values.

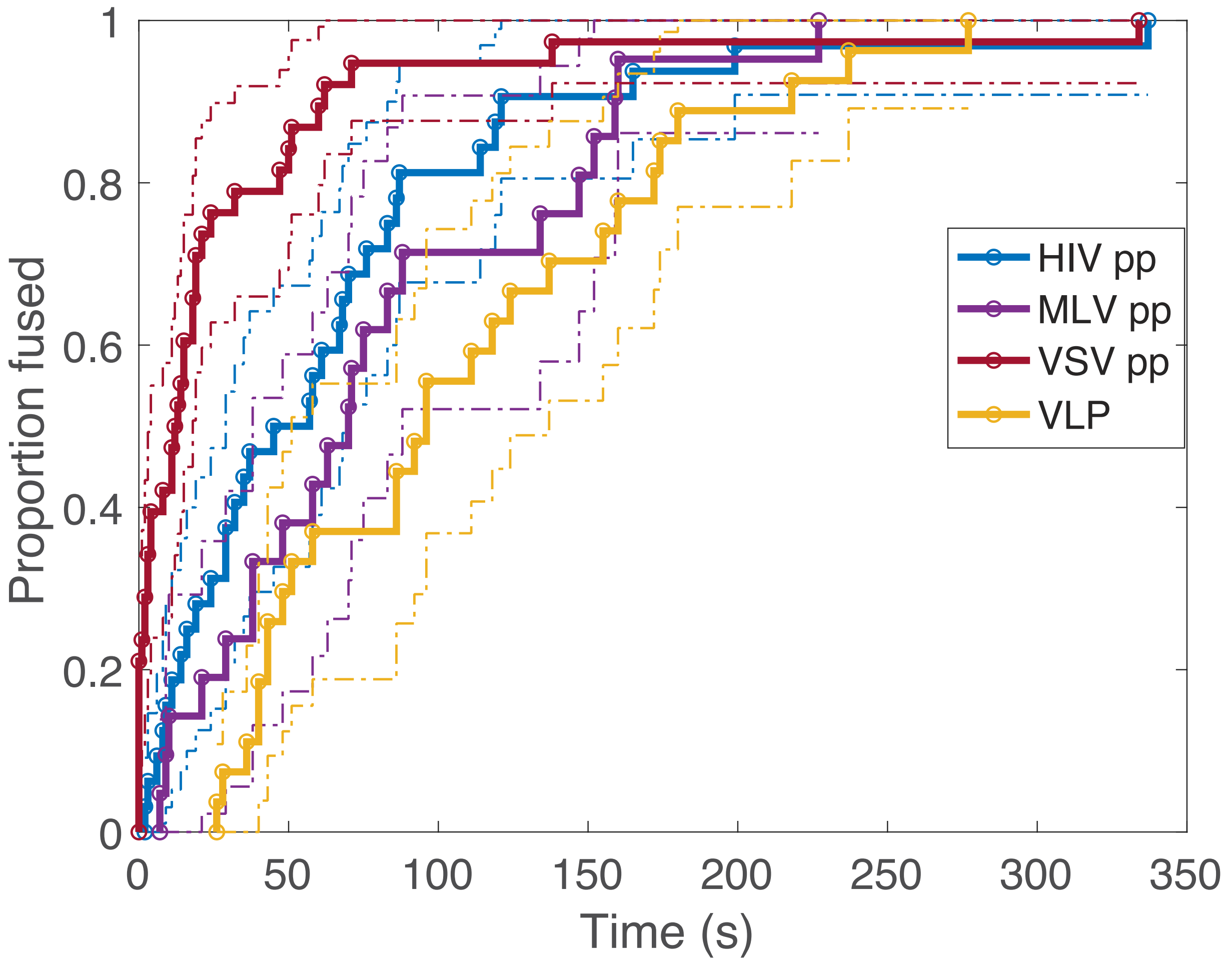


**Figure S8. Single-virus lipid mixing using virus-like particles is not significantly different from HIV or MLV pseudovirions**. Single-virus lipid mixing measurements were made using virus-like particles (VLPs) in an assay otherwise identical to the pseudovirus lipid-mixing assay. Cumulative distribution functions were calculated using 27 lipid-mixing events and compared to the previously measured HIV, MLV, and VSV pseudovirus distributions. VLP lipid mixing was slightly but not significantly slower than HIV and MLV lipid mixing although significantly slower than VSV ($p < 10^{-6}$, Kolmogorov-Smirnov test with Bonferroni multiple-hypothesis correction).

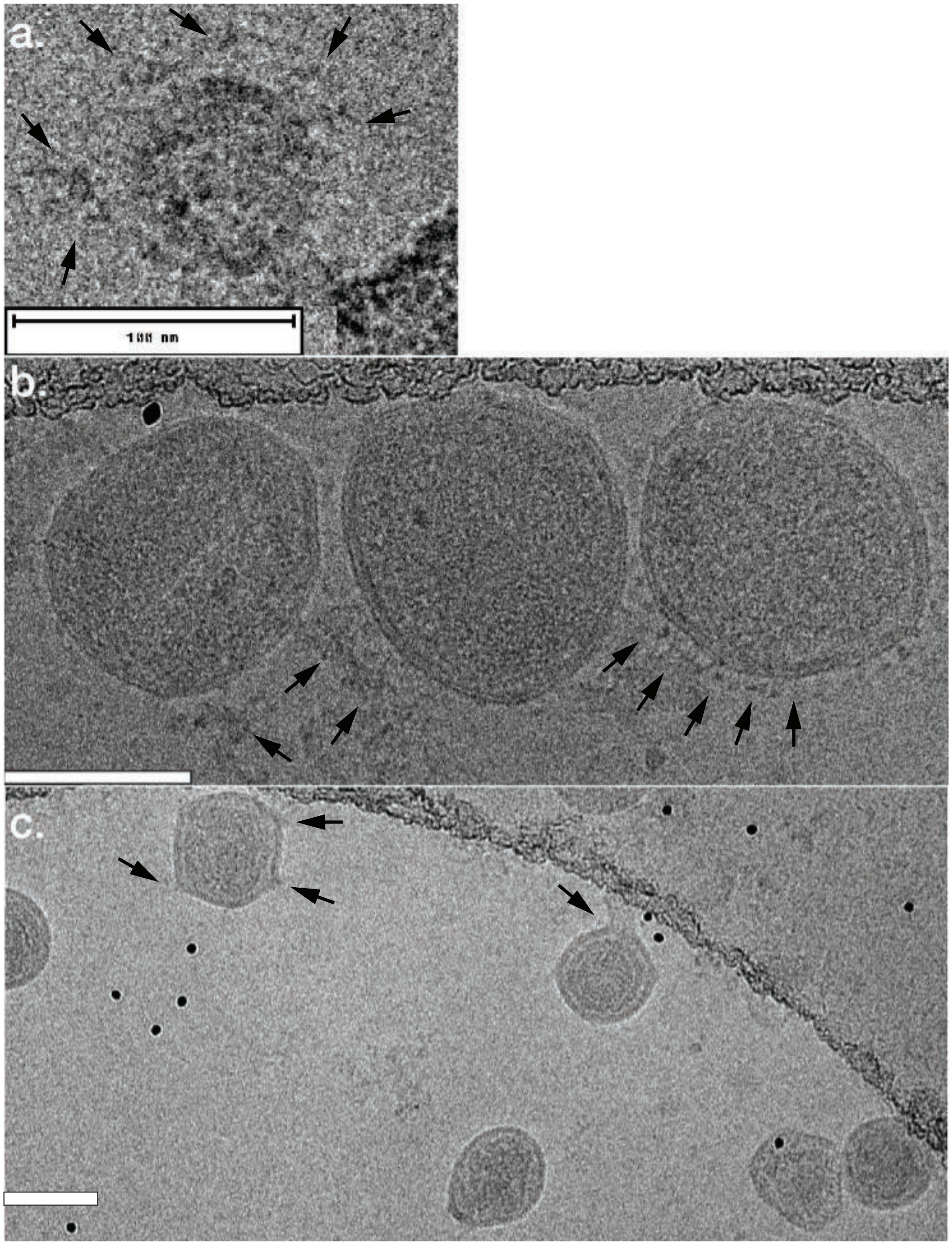


**Figure S9**. Electron cryo-micrographs of VSV (panel a), HIV (panel (b), and MLV pseudoviral particles (panel c). Scale bars denote 100 nm. Spike density varies but in general fewer SARS-CoV-2 spikes were visualized on MLV pseudoviral particles than on VSV and HIV particles. Arrows denote density putatively assigned to spike glycoproteins.

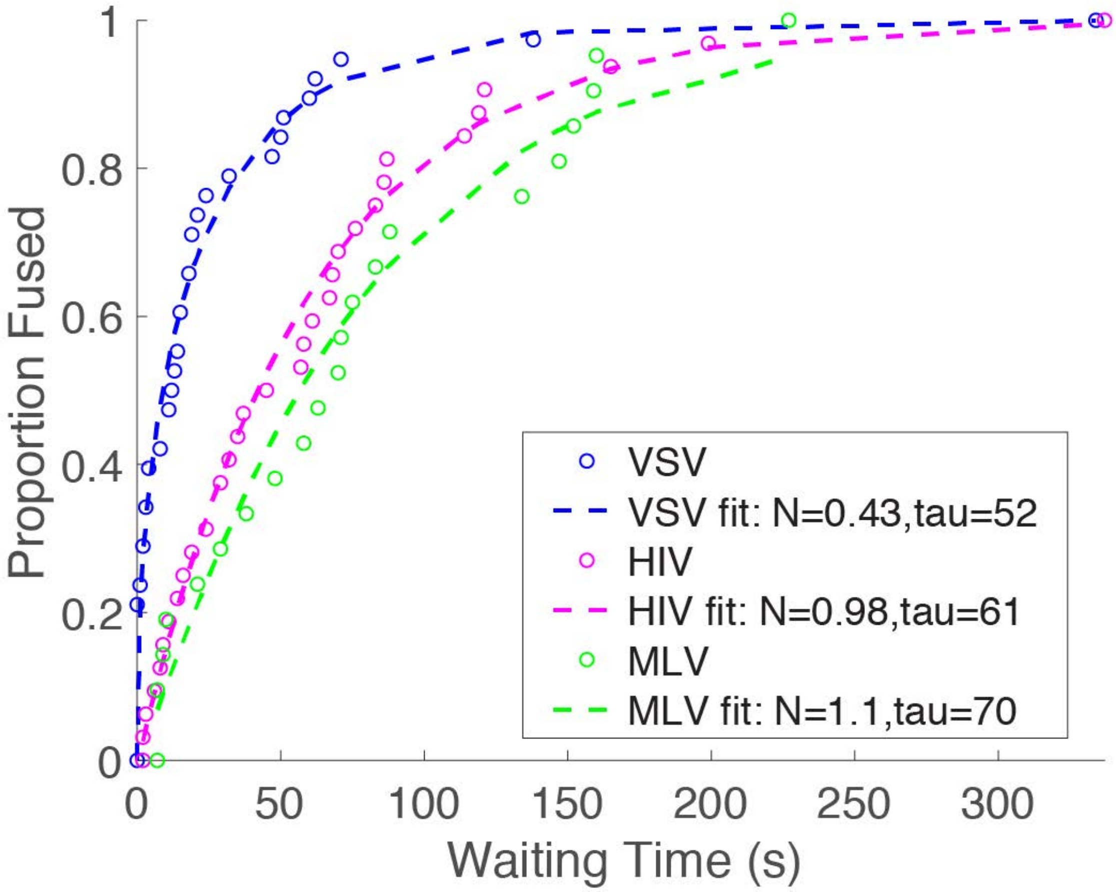


**Figure S10. Gamma distribution fits support a model where different pseudovirus backgrounds have the same fusion stoichiometry but different rates**. Plotted here are independent gamma-distribution fits to each cumulative distribution function. Based on Akaike Information Criteria, the agreement is not sufficiently better to merit the additional model parameters (AIC 2530 for fits shown here versus 2470 for fits shown in the main text). A time lag was tested as an additional fit parameter in all fits, but the maximum-likelihood value was less than 1s in all cases.

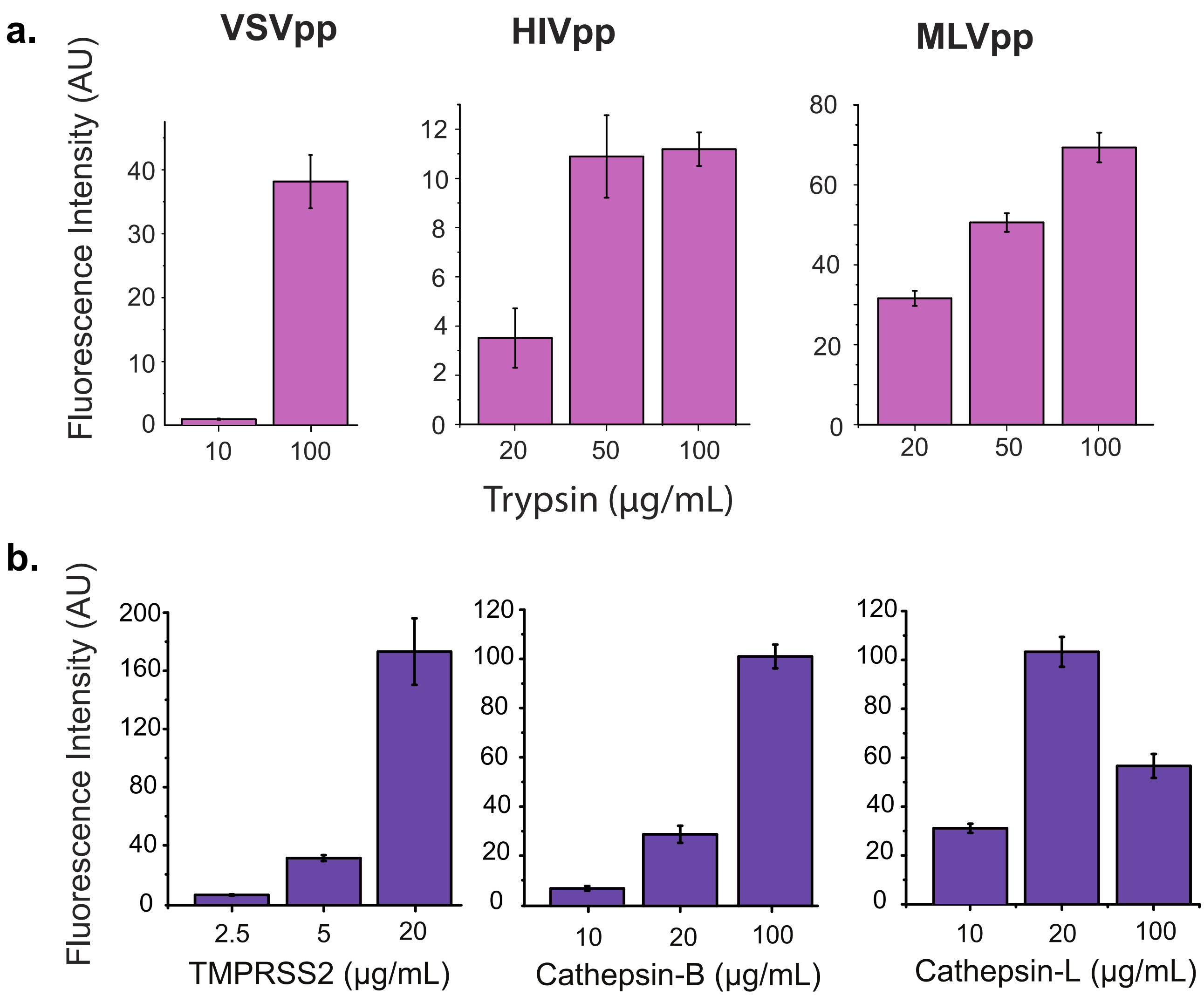


**Figure S11.** Mean fluorescence intensity change over all pixels in a 134x134 µm field of view measured from start to end of fusion for different pseudoviruses with trypsin treatment at different concentrations (top panel) and MLV pseudoviruses, treated with different proteases (bottom panel). This yields a bulk estimate of lipid mixing. Values are plotted as averages over two fields of view from independent experiments. Error bars represent 90% confidence intervals calculated via bootstrap resampling performed by dividing each field of view into 9 equal tiles.

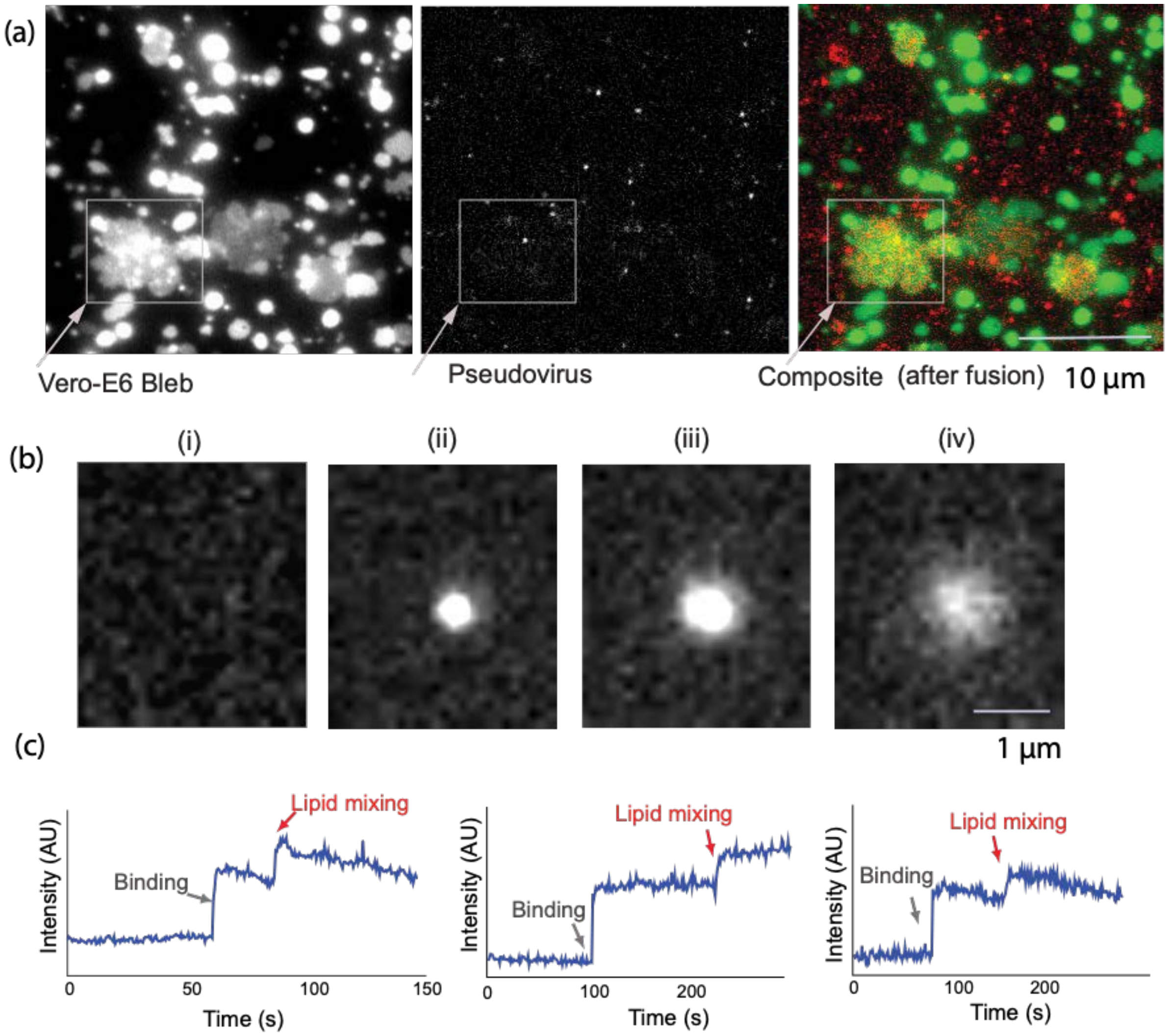


**Figure S12.** Fusion of SARS-CoV-2 pseudovirus activated with TMPRSS11D (Human Airway Trypsin-like protease or HAT) with Vero-E6 plasma membrane vesicles (blebs). Cells were labeled with DiO membrane dye to facilitate visualization of vesicles, and pseudoviruses were labeled with Texas Red membrane dye. Panel (a) shows fluorescence images of labeled vesicles, dequenching of Texas Red dye of pseudovirus after fusion, and an overlay showing colocalization of Texas Red dispersion onto plasma membrane vesicles. Panel (b) shows a single spot in the Texas Red channel prior to binding (i), at time of binding (ii), and at time of lipid mixing (iii). Panel (c) shows representative fusion traces, plotted as the total intensity of a single-virus spot.
